# Antiphage defence systems Druantia III and Zorya II synergise via shared DNA intermediates in a phage-specific manner

**DOI:** 10.64898/2026.05.12.724681

**Authors:** Yi Wu, Zhiying Zhang, Sofya K. Garushyants, Peter R. Weigele, Eugene V. Koonin, Dinshaw J Patel, Franklin L. Nobrega

## Abstract

Most bacteria encode multiple antiphage defence systems, but how these systems interact, remains poorly understood. Here, we define the mechanism of Druantia III and explore its synergy with Zorya II. Druantia III is a late-acting defence where DruH is the likely infection sensor and DruE is a helicase-nuclease effector that engages ssDNA-containing replication intermediates. Druantia III and Zorya II are each sensitive to the loss of RecD, whereas their synergy requires an intact RecBCD complex, indicating that the combined response depends on a shared DNA-processing hub. During T3 infection, Zorya apparently preserves this hub by limiting accumulation of the phage RecBCD-inhibitor Gp5.9, whereas DruE together with RecBCD promote the formation of DNA structures permissive for ZorE cleavage. During Bas37 infection, Druantia provides the dominant pathway and recruits ZorE in a non-canonical, ZorAB-independent manner. These findings show how shared DNA intermediates can connect defence systems into a coordinated antiphage response.

## Introduction

Bacterial antiphage defence is often described one system at a time, but most bacteria encode multiple defence systems in the same genome^1–3^. Comparative genomic analyses have shown that these systems do not assort randomly^1,3,4^, and experimental work is beginning to explain why. Some combinations broaden protection by acting in parallel against different phages or different DNA chemistries, for example, in defence islands that combine BREX with type IV restriction^5,6^. Others are hierarchically coupled, with one system creating molecular substrates for another, exemplified by restriction cleavage stimulating spacer acquisition by CRISPR-Cas systems^7^. Still others act as safeguards or layered responses, in which loss, sabotage, or bypass of one defence system exposes or activates another^8–11^. Finally, there are cases of genuine synergy, in which protection emerges or is greatly enhanced only from the combination of two systems^4,12^. Together, these findings suggest that bacterial immunity is shaped not only by the activities of individual defence systems, but also by multiple types of interactions between those.

One such interaction is the synergy between Druantia type III and Zorya type II. This pair was recently identified in *Escherichia coli* as a robust example of defence-system synergy, but the mechanism behind it remained unknown^4^. Zorya that was first identified in defence-island surveys is built around the conserved membrane proteins ZorA and ZorB, with various soluble components defining distinct subtypes^13^. Recent structural and biochemical work showed that the cores of Zorya type I and II form a 5:2 inner-membrane complex with features characteristic of ion-driven rotary motors^14,15^. In type II Zorya systems, this complex is proposed to recruit the soluble effector ZorE, a nickase required for population-wide immunity^15^. These findings provide a mechanistic framework for how Zorya links phage-triggered membrane-associated events to downstream DNA damage.

By contrast, the mechanism of action of Druantia remained unresolved. There are several subtypes of Druantia that all share the DruE protein but differ in their associated genes^13^. Druantia III is the most compact variant of this system consisting of only two components, DruE and DruH, whereas other subtypes include additional genes. Sequence analysis predicted that DruE is a helicase, whereas DruH remained uncharacterised^13^. Recent comparative analysis placed DruE within the YprA-like family of defence-associated SF2 (superfamily 2) helicases. The YprA-like family spans components of various defence systems, such as DISARM (Defense Island System Associated with Restriction-Modification), Dpd (7-deazapurine in DNA), ARMADA (disARM-related Antiviral Defense Array) and Druantia, in which helicases are often coupled with C-terminal nucleases^16^. These observations place Druantia III in a larger class of related DNA-processing immune systems, but do not explain how this system senses infection, what is the exact mechanism of defence, or how Druantia III cooperates with Zorya II.

Here, we present a mechanistic model for Druantia III and use it to explain the synergy between Druantia III and Zorya II. We show that Druantia III is a late-acting defence system in which DruH is likely the phage infection sensor and DruE is a helicase-nuclease effector. Our findings support a model in which DruH is activated by late phage proteins and acts on late infection-associated DNA structures, possibly generating or exposing DNA states that can be productively engaged by DruE. We further show that the synergy between Druantia III and Zorya II depends on distinct phage-dependent routes linked by DNA intermediates and their accessibility. Druantia III and Zorya II are each sensitive to the loss of RecD, whereas their synergy requires an intact RecBCD complex, indicating that the combined response depends on a shared DNA-processing hub. During T3 infection, Druantia alone does not provide detectable protection, but the escape mutant data indicate that it can still sense infection, whereas its downstream contribution to defence is normally constrained. Zorya safeguards the RecBCD-dependent synergistic state by acting early enough to limit expression of the phage-encoded RecBCD inhibitor Gp5.9, whereas DruE together with RecBCD likely helps maintain DNA states permissive for ZorE activity. During Bas37 infection, Druantia provides the dominant defence pathway and appears to promote a related ZorE-permissive DNA state in a non-canonical, ZorAB-independent manner, through a downstream RecBCD-linked DNA-processing step and, potentially, by limiting RecA occupancy of shared ssDNA-rich intermediates. Together, these findings show how shared DNA intermediates can connect defence systems into coordinated antiviral responses.

## Results

### Recurrent functional association between Druantia III and Zorya II

Druantia III and Zorya II synergise against distinct groups of phages, including *Tequatrovirus* and *Autographiviridae* (**Figure 1A**)^4^. To define how each system contributes to this response, we tested versions lacking individual genes (**Figure 1B**). The results showed that which proteins are required for synergy, depends on the targeted phage. In the case of Bas37, a phage targeted by Druantia III but not Zorya II alone, DruE and DruH together with ZorE were sufficient to reproduce the full combined phenotype. By contrast, in the case of T3, a phage targeted by Zorya II but not Druantia III, the full Zorya II and Druantia III machinery were required.

**Fig. 1.**
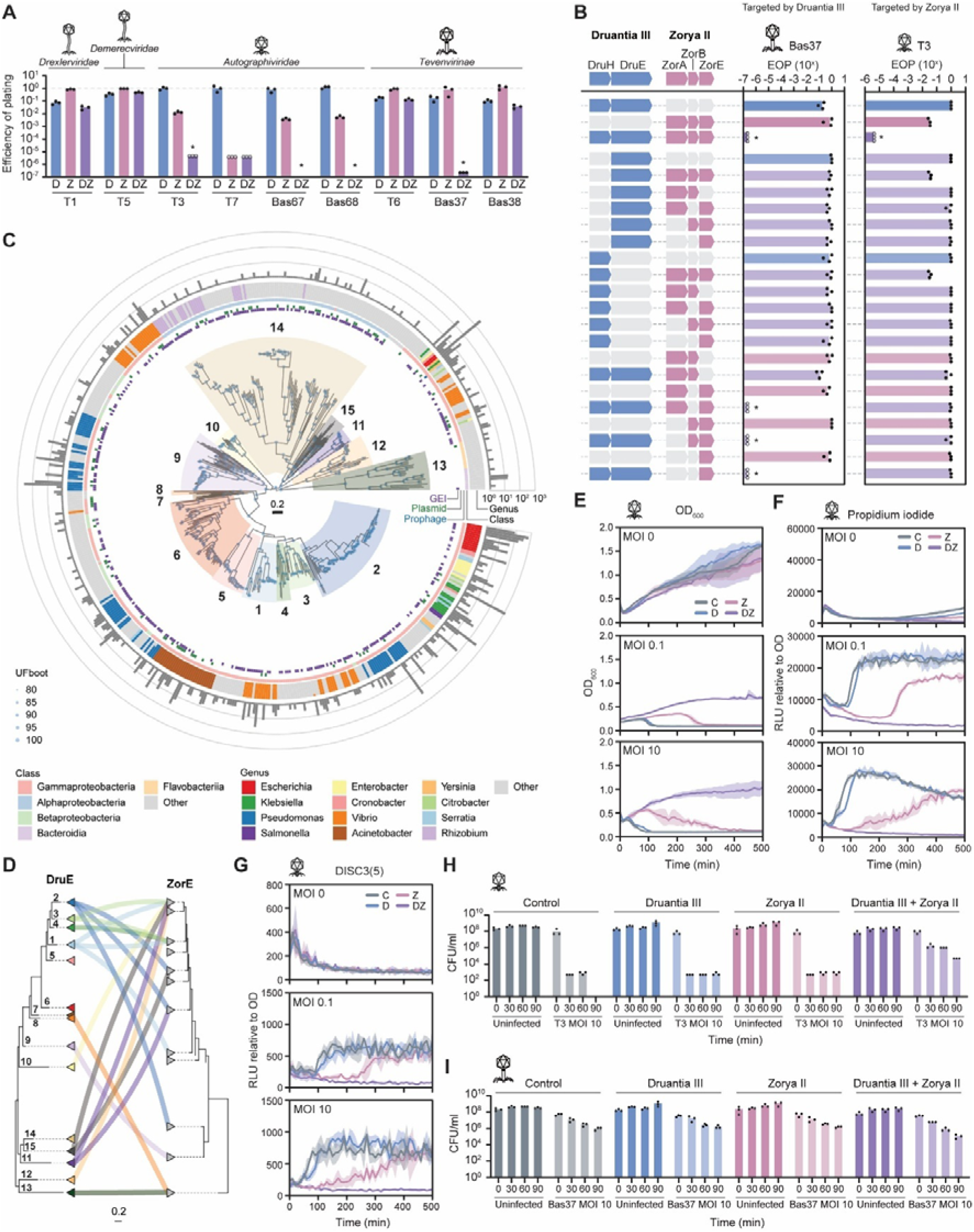
Recurrent association and phage-dependent synergy between Druantia III and Zorya II. **(A)** Efficiency of plating (EOP) assays for Druantia III (D), Zorya II (Z), or both systems (DZ) against the indicated *Drexlerviridae*, *Demerecviridae*, *Autographiviridae* and *Tevenvirinae* phages. Bars represent mean, with dots indicating individual replicates. Asterisks indicate statistically significant (p < 0.05) differences between DZ and the individual defences. **(B)** Genetic analysis of the components required for synergy. Left, schematics indicate the presence or absence of Druantia III genes (*druH* and *druE*) and Zorya II genes (*zorA*, *zorB* and *zorE*) in the tested constructs. Right, EOP values for Bas37, a phage targeted by Druantia III, and T3, a phage targeted by Zorya II. Bars represent mean, with dots indicating individual replicates. Asterisks indicate statistically significant (p < 0.05) differences between DZ and the individual defences. **(C)** Maximum-likelihood phylogeny of dereplicated DruE proteins from complete Druantia III systems. The 15 DruE clusters are indicated. Node support is shown by ultrafast bootstrap (UFboot) values, and the outer tracks indicate taxonomic assignment and genomic context, including association with genomic islands (GEIs), plasmids and prophages. **(D)** Tanglegram linking DruE clusters to co-occurring ZorE clades, showing that closely related Druantia III variants can associate with distinct Zorya II lineages. The full tanglegram with uncollapsed branches is shown in Figure S1D. **(E-G)** Population-level response to T3 infection in control cells (C) and cells expressing Druantia III (D), Zorya II (Z), or both systems (DZ). Bacterial growth was measured as optical density at 600 nm (OD_600_) (E), membrane damage as propidium iodide uptake normalised to OD (F), and membrane depolarisation as DISC3(5) fluorescence normalised to OD (G), at the indicated multiplicities of infection (MOI). Lines and shaded areas indicate mean ± standard deviation. **(H,I)** Time-course quantification of viable cells (CFU/ml) during infection with T3 (H) or Bas37 (I) at MOI of 10 in control cells and cells expressing Druantia III, Zorya II, or both systems. Bars represent mean with dots indicating individual replicates.

We next explored the distributions of Druantia III and Zorya II across bacterial, to determine whether the functional coupling between these defence systems reflected a broad association or was restricted only to the specific model systems studied here. Using precomputed PADLOC annotations for RefSeq version 209^17–19^, we identified 6,885 bacterial genomes encoding complete Druantia III systems, defined by the presence of both DruE and DruH. These systems were found predominantly in Proteobacteria, especially Gammaproteobacteria (**Table S1**). After dereplication, the dataset included 555 unique DruE proteins and 577 unique DruH proteins. Phylogenetic reconstructions for both protein families revealed strongly congruent topologies, with little evidence of mixing and matching between DruE and DruH variants (**Figure 1C**, **Figure S1A,B**). This pattern suggests that Druantia III typically spreads as an intact module and/or that specific DruE-DruH pairings are maintained by functional constraints. The Druantia III system characterised here belongs to cluster 11, a clade detected predominantly in *Escherichia* (4,288 genomes) and *Klebsiella* (246 genomes), and frequently carried on integrated SPIDER phage-like elements together with other defence systems, such as Zorya II and ARMADA II^16^. However, the overall diversity of Druantia III is much wider, with 15 distinct clades identified in DruE and DruH phylogenetic trees and associated with specific bacterial lineages and habitats, indicating that this system is not restricted to a single lineage or ecological niche (**Figure 1C**, **Figure S1A**; habitat annotations are provided in the associated dataset, see Key Resources Table). Thus, our experimental work focused on one representative branch of the widespread and evolutionarily diverse Druantia III family.

We next asked whether the association between Druantia III and Zorya II extends beyond cluster 11 studied here. To address this question, we reconstructed a phylogeny of ZorE proteins from complete Zorya II systems in RefSeq version 209^17^ and retained only variants containing a recognisable endonuclease domain. We identified 4,156 genomes encoding this form of ZorE and, after dereplication, obtained 433 unique ZorE sequences. ZorE variants co-occurring with Druantia III were distributed across multiple branches of the ZorE phylogeny (**Figure 1D, Figure S1C,D**). Although members of Druantia III cluster 11 most often co-occurred with ZorE from one major branch, this association was not exclusive, and ZorE proteins from more distant branches were also found linked to closely related Druantia III variants. Overall, ZorE co-occurred with 11 Druantia III clusters and was frequently located within the same defence island, often within a 20-kb genomic neighbourhood (**Figure S1D**). These observations indicate that the association between Druantia III and Zorya II is not confined to a single class of mobile genetic element (such as SPIDERs) or one specific variant combination but instead represents a broader and recurrent linkage that likely reflects widespread synergy. We also inspected the ZorE proteins from clusters that co-occur with Druantia III and compared these to those from branches devoid of Druantia III. The majority of Druantia-associated ZorE proteins (53 of the 59 leaves in the ZorE tree, 90%) contained an additional C-terminal region with a nuclease-like signature of conserved amino acid residues. Although this region could not be confidently assigned to a defined nuclease family, it contains a cluster of conserved residues that group together in the predicted structure to form a pocket-like surface, including acidic and basic residues reminiscent of the catalytic architecture found in many nucleases, such as the PD-(D/E)xK superfamily (**Figure S1E**). The recurrent presence of this domain among the Druantia-associated variants of ZorE therefore suggests that this extension is a conserved feature of the synergy-competent ZorE subset (**Figure S1D**).

Analysis of the genomic neighbourhood surrounding Druantia III loci further supported that, rather than occurring in isolation, Druantia III was commonly embedded within defence-rich regions that also encoded other DNA-targeting systems. In several Druantia III clusters, the surrounding 20-kb regions were enriched in type I and type IV restriction-modification (RM) systems (**Figure S1F**, **Table S2**). Thus, Druantia III is frequently integrated into a broader class of defence islands, where it might act alongside other systems that monitor or process foreign DNA. Together, the genetic dissection and comparative genomic analysis support the view that Druantia III and Zorya II are recurrently associated defence modules whose cooperation is biologically relevant and conserved across diverse bacterial backgrounds.

### Combination of Druantia III and Zorya II can shift defence towards survival of the infected cell

We next asked whether the Druantia III and Zorya II individually and their combination enabled survival of the infected cells or instead produce a defence outcome associated with the death of the infected cells^20^. To distinguish between these possibilities, we monitored bacterial growth in the absence and in the presence of phage at both high and low multiplicities of infection (MOI) (**Figure 1E**), while simultaneously measuring propidium iodide uptake as a readout of membrane damage (**Figure 1F**), and DISC_3_(5) fluorescence as a readout of membrane depolarisation (**Figure 1G**). When challenged with phage T3, Zorya II alone provided protection via a mechanism that stalled cell growth, and increased membrane damage and membrane depolarisation at high phage MOI, indicating that infected cells did not survive under these conditions. By contrast, when Zorya II was combined with Druantia III, the bacterial population remained largely protected and showed little or no evidence of membrane damage or depolarisation under the same conditions.

To assess this phenotype more directly, we followed viable cell counts over time during liquid infection assays performed at high MOI (**Figure 1H**). Consistent with the physiological measurements above, the Druantia III-Zorya II combination yielded markedly greater cell survival rates than either system alone (which behaved similarly to the control) during T3 infection. In contrast, under Bas37 infection, Druantia III alone did not preserve cell viability relative to the control, and the combination with Zorya led to an even greater depletion of viable cells (**Figure 1I**). Thus, combining the two systems enhanced the defence response against Bas37, but with a cellular outcome clearly distinct from that observed against T3. Together, these results show that, under certain conditions, as in the case of the T3 infection, synergy between Druantia III and Zorya II can shift defence from a state in which infected cells are severely compromised to one that permits survival and recovery.

### Synergy between Druantia III and Zorya II depends on RecBCD

Because the activities of both Zorya II and Druantia III involve interactions with DNA^13–15,21^, we asked whether DNA mimic proteins could alter the activity of either system alone or affect the synergistic phenotype. To address this question, we tested a panel of DNA mimic proteins against each defence system individually and in combination. Several of these proteins affected defence, but with clearly distinct outcomes (**Figure 2A**). For Zorya II, DinI enhanced protection against T3. At high concentrations, DinI inhibits RecA binding to ssDNA and therefore stalls the SOS response^22^, suggesting that the activity of Zorya II could be stimulated by prevention of DNA repair processes that might repair the phage genome or by the presence of RecA-unbound, free ssDNA.

**Figure 2.**
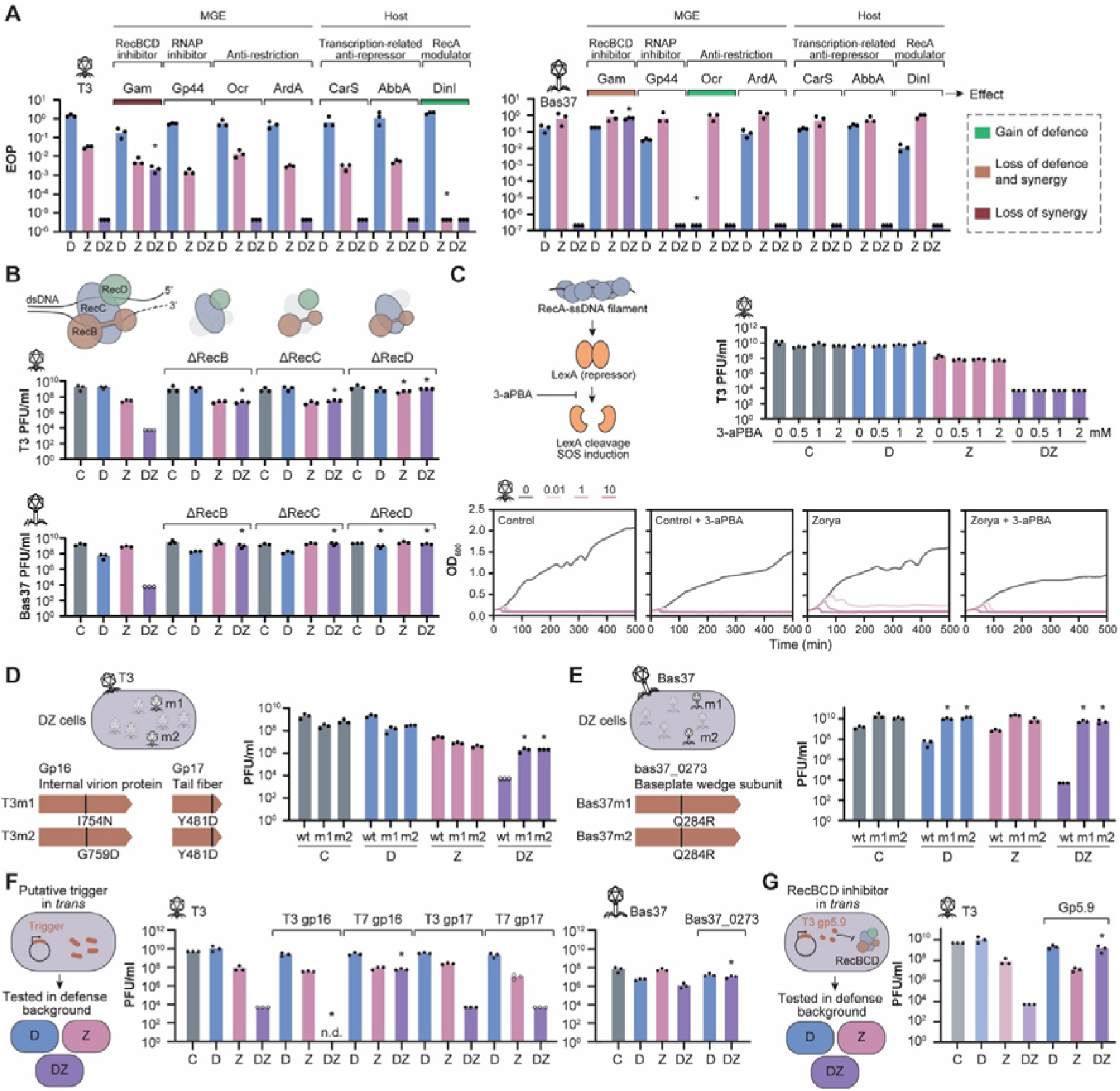
DNA mimic proteins, RecBCD dependence, and phage triggers reveal distinct routes into Druantia III-Zorya II synergy. **(A)** Efficiency of plating (EOP) of phages T3, T7, and Bas37 on cells carrying Druantia III alone (D), Zorya II alone (Z), or both systems together (DZ), in the presence of the indicated phage-encoded or host-derived DNA mimic proteins expressed *in trans*. Tested proteins are grouped by annotated function. Colours indicate the qualitative outcome of each perturbation: gain of defence, loss of defence and synergy, or loss of synergy. Asterisks indicate statistically significant differences to the same cells without a DNA mimic. Bars represent mean, with dots indicating individual replicates. **(B)** Phage infection assays in strains lacking individual RecBCD subunits. T3 and Bas37 plating was measured on control cells (C) and on cells carrying Druantia III (D), Zorya II (Z), or both systems together (DZ) in Δ*recB*, Δ*recC*, and Δ*recD* backgrounds. Asterisks indicate statistically significant differences to the same cells with intact RecBCD. Bars represent mean, with dots indicating individual replicates. **(C)** Effect of 3-aPBA on Zorya II defence against phage T3. Left, schematic of the assay rationale. 3-aPBA inhibits LexA self-cleavage and thereby blocks SOS induction downstream of RecA-ssDNA filament formation. Top right, T3 titres measured in C, D, Z, and DZ cells in the presence of the indicated concentrations of 3-aPBA. Bars represent mean, with dots indicating individual replicates. Bottom, OD_600_ of control and Zorya II-expressing cells in the absence or presence of 1 mM 3-aPBA during T3 infection at the indicated multiplicities of infection (MOI). **(D)** Isolation and characterisation of T3 escape mutants recovered on DZ cells. Left, schematics of the two independent escaper mutants and their mutations in *gp16* (internal virion protein) and *gp17* (tail fibre). Right, plating of wild-type (wt) T3 and the escaper mutants on C, D, Z, and DZ cells. Asterisks indicate statistically significant differences between the escaper and wt phages in the same defence background. Bars represent mean, with dots indicating individual replicates. **(E)** Isolation and characterisation of Bas37 escape mutants recovered on DZ cells. Left, schematics of the two independent escaper mutants, both carrying the same Q284R substitution in *bas37_0273*, encoding a predicted baseplate wedge subunit. Right, plating of wild-type Bas37 and the escaper mutants on C, D, Z, and DZ cells. Asterisks indicate statistically significant differences between escaper and wt phages in the same defence background. Bars represent mean, with dots indicating individual replicates. **(F)** Effect of expressing putative phage triggers in *trans* on phage infection in the indicated defence backgrounds. Left, schematic of the assay design. Middle, T3 plating on control and defence-system-containing cells in the absence or presence of T3 gp16, T7 gp16, T3 gp17, or T7 gp17 expressed *in trans*. Right, Bas37 plating on control and defence-system-containing cells in the absence or presence of Bas37_0273 expressed *in trans*. Asterisks indicate statistically significant differences to the same cells in the absence of putative phage trigger; n.d., not detected. Bars represent mean, with dots indicating individual replicates. **(G)** Effect of expressing T3 Gp5.9, a RecBCD inhibitor, *in trans* on T3 infection in the indicated defence backgrounds. Left, schematic of the assay design. Right, T3 plating on control and defence-system-containing cells in the absence or presence of Gp5.9 expressed *in trans*. Asterisks indicate statistically significant differences relative to the same cells without Gp5.9. Bars represent mean, with dots indicating individual replicates.

The activity of Druantia III against Bas37 was enhanced by Ocr (**Figure 2A**), but co-immunoprecipitation of Ocr with either DruE or DruH failed to identify direct interactions between the proteins (**Figure S2A-D**). This finding suggests that Ocr might enhance Druantia III defence through indirect mechanisms, such as inhibition of other host defence pathways, given that Ocr has been shown to interact with type I restriction-modification enzymes and with the BREX methyltransferase BrxX/PglX^23,24^.

Unexpectedly, Gam abolished the synergy against T3 and Bas37, without affecting the activity of Druantia III or Zorya II individually (**Figure 2A**). Therefore, we hypothesised that the inhibition of synergy was not caused by a direct interaction between Gam and the defence system proteins but rather reflected interference with a pathway required for the combined response. Consistent with this possibility, we found no evidence that Gam directly interacted with the defence proteins themselves using pull-down assays (**Figure S2E-G**). Because Gam is a known inhibitor of RecBCD^25^, the major host helicase-nuclease complex involved in the processing of double-stranded DNA ends^26^, we asked whether the Gam phenotype could be explained by RecBCD inhibition. We tested defence in *recBCD* mutant backgrounds^27^ and observed that both Druantia III and Zorya II individually were sensitive to the loss of *recD*, but not of *recB* or *recC* (**Figure 2B**). RecBCD normally loads RecA onto 3’ ssDNA in a Chi-dependent manner, whereas RecBC lacking RecD loads RecA constitutively in a Chi-independent manner, which is expected to reduce the pool of accessible ssDNA-containing intermediates^28,29^. Together with the findings on DinI, these observations in a Δ*recD* background reinforce the hypothesis that the activity of Zorya II depends either on DNA intermediates containing accessible ssDNA regions free of RecA, or on the lack of downstream SOS response activation. To test this hypothesis and distinguish between these two possibilities, we used 3-aPBA, an inhibitor of LexA self-cleavage, to block the SOS response downstream of RecA loading onto ssDNA^30,31^. If the SOS response were inhibitory, Zorya II activity would be expected to increase under these conditions. Instead, we observed that 3a-PBA had no effect on Zorya II defence in EOP assays and appeared to decrease it in liquid assays (**Figure 2C**). Together with the DinI and Δ*recD* results, these findings do not support downstream SOS induction as the main component of the inhibitory effect and are better consistent with RecA occupancy of ssDNA being the more relevant factor affecting Zorya II activity.

By contrast, the behaviour of the combined system pointed to a distinct requirement at the level of synergy. Although the activities of Druantia III and Zorya II individually were not affected by deletion of *recB* or *recC*, synergy was impaired in each *recBCD* mutant background, indicating that an intact RecBCD complex is specifically required in the synergy context (**Figure 2B**).

Bas37 is a T4-like phage and encodes a gp2-related protein (Bas37_0266) that is predicted to protect the ejected phage DNA ends from RecBCD, as described for T4 gp2^32,33^. Nevertheless, Druantia III-Zorya II synergy against Bas37 was still observed (**Figure 1A**). This observation indicates that the RecBCD-dependent step required for synergy is unlikely to be a simple attack on the DNA ends of the incoming phage. Consistent with this possibility, Gam, which blocks RecBCD by occupying its DNA-binding surface^25^, abolished synergy, whereas gp2-like protection of the injected genome did not. Together, these results indicate that synergy depends on a RecBCD-mediated DNA-processing step beyond direct targeting of the initially injected phage genome.

Further support for this connection came from comparative genomic analysis. Using HMM^34,35^ profiles to search for RecBCD components across representative Druantia III-containing genomes, we found that RecC, the subunit unique to RecBCD, was broadly present across Druantia III clades and in essentially all representatives in which Druantia III co-occurred with Zorya II, with only rare exceptions in a divergent clade (**Figure S2H**). Thus, both the genetic data and the comparative distribution are consistent with intact RecBCD acting as a recurrent DNA-processing hub for Druantia III-Zorya II synergy.

### Late phage triggers and RecBCD inhibition define distinct routes to Druantia-Zorya synergy

To understand how Druantia III and Zorya II sense infection in a synergistic context, we isolated T3 and Bas37 escape mutants from cells expressing both systems. Surprisingly, the T3 escape mutants remained susceptible to Zorya II alone, but escaped the additional protection conferred by the combined systems (**Figure 2D**), suggesting that the mutations allowed the phage to escape the Druantia III branch of synergy. Bas37 mutants escaped individual defence by Druantia III and the combined Druantia III-Zorya II defence (**Figure 2E**), indicating that these mutations also ensured escape from the Druantia III branch. Sequencing of the T3 and Bas37 escapers revealed mutations in structural phage proteins expressed late in the phage life cycle, namely, tail fibre gp17 (Y481D in both mutants) and internal virion protein gp16 (I754N in T3m1 and G759D in T3m2) in T3, and baseplate wedge subunit bas37_0273 (Q284R in both mutants) in Bas37, suggesting that Druantia III provides protection at the late stage of infection.

To test whether these phage proteins were directly sensed by Druantia III, we expressed them *in trans* and assessed their effects on defence. Expression of T3 gp16, but not gp17, enhanced synergy against T3 without enhancing the Zorya response, suggesting activation of the Druantia branch during synergy (**Figure 2F**). By contrast, the homologous T7 gp16 did not trigger defence and instead inhibited synergy. T3 gp16 and T7 gp16 differ at only 15 of 1318 residues, indicating that limited sequence variation is sufficient to convert an activating phage protein into a non-activating competitor. The ability of T7 gp16 to inhibit synergy when expressed *in trans* is therefore consistent with retention of sensor binding despite escape from Druantia activation. For Bas37_0273, expression *in trans* did not detectably activate Druantia III (**Figure 2F**). However, co-immunoprecipitation detected a weak association between Bas37_0273 and DruH, whereas the escape variant Bas37_0273^Q284R^ showed no detectable interaction, consistent with weak or transient engagement of DruH by the wild-type protein (**Figure S2H**). Because Bas37_0273 is involved in baseplate assembly, it is possible that Druantia III recognises Bas37_0273 only in the context of a higher-order phage structure, such that the isolated protein is insufficient for productive engagement and activation.

Taken together, these results indicated that T3 activates both Zorya II and the Druantia III branch of the combined response, but that synergy requires an additional RecBCD-dependent step. T3 encodes Gp5.9, a direct inhibitor of RecBCD that blocks access to the DNA-binding surface of the complex^36^. We therefore hypothesised that Gp5.9 supresses the RecBCD-dependent DNA-processing state required for Druantia III-Zorya II synergy. In the context of Druantia III-Zorya II synergy, given that Zorya II acts early during infection^14,15^, it could limit the accumulation of Gp5.9 and thereby preserve RecBCD activity, allowing the combined response to proceed. To test this hypothesis, we expressed Gp5.9 *in trans*, reasoning that its pre-expression would inhibit RecBCD regardless of the timing of phage infection and therefore suppress synergy. Expression of Gp5.9 *in trans* indeed inhibited synergy, while not abolishing Zorya II activity (**Figure 2G**).

These results indicate that phage-encoded inhibition of RecBCD can specifically block the RecBCD-dependent step required for Druantia III-Zorya II synergy, even when infection is still sensed.

In summary, these results suggest that synergy between Druantia III and Zorya II is achieved via two distinct routes. Against T3, Zorya acts upstream of Druantia III, early enough to preserve the RecBCD-dependent DNA-processing state required for synergy, likely by limiting the accumulation of RecBCD-inhibiting phage proteins. By contrast, against Bas37, Druantia III acts upstream of Zorya II, sensing phage infection (the sensing Zorya components ZorAB are dispensable for synergy) and likely co-opting ZorE for additional effector activity.

### DruE is a YprA-like family helicase that preferentially engages 3’-tailed DNA

To better understand the defence mediated by Druantia III and therefore its synergy with Zorya II, we set out to characterise the molecular mechanism of Druantia III. To this end, we first focused on DruE, the only protein shared across all known Druantia subtypes^13^ and therefore the most plausible candidate for a conserved core activity. Recent comparative analysis placed DruE within the YprA-like family of defence-associated SF2 helicases, indicating that Druantia III belongs to a broader class of immune systems centred around helicases of this family^16^. Sequence analysis indicated that DruE contains an N-terminal P-loop ATPase region characteristic of SF1/SF2 helicases, including the DEAH signature of SF2, a conserved C-terminal helicase-associated region, and an MrfA-like Zn-binding module (**Figure 3A**). Additionally, downstream of the helicase, DruE contains two C-terminal phospholipase D (PLD) superfamily nuclease domains, including an upstream domain predicted to be inactive (PLDi) and a downstream domain that retains the features of an active PLD module (PLDa) (**Figure 3A**)^16^.

**Figure 3.**
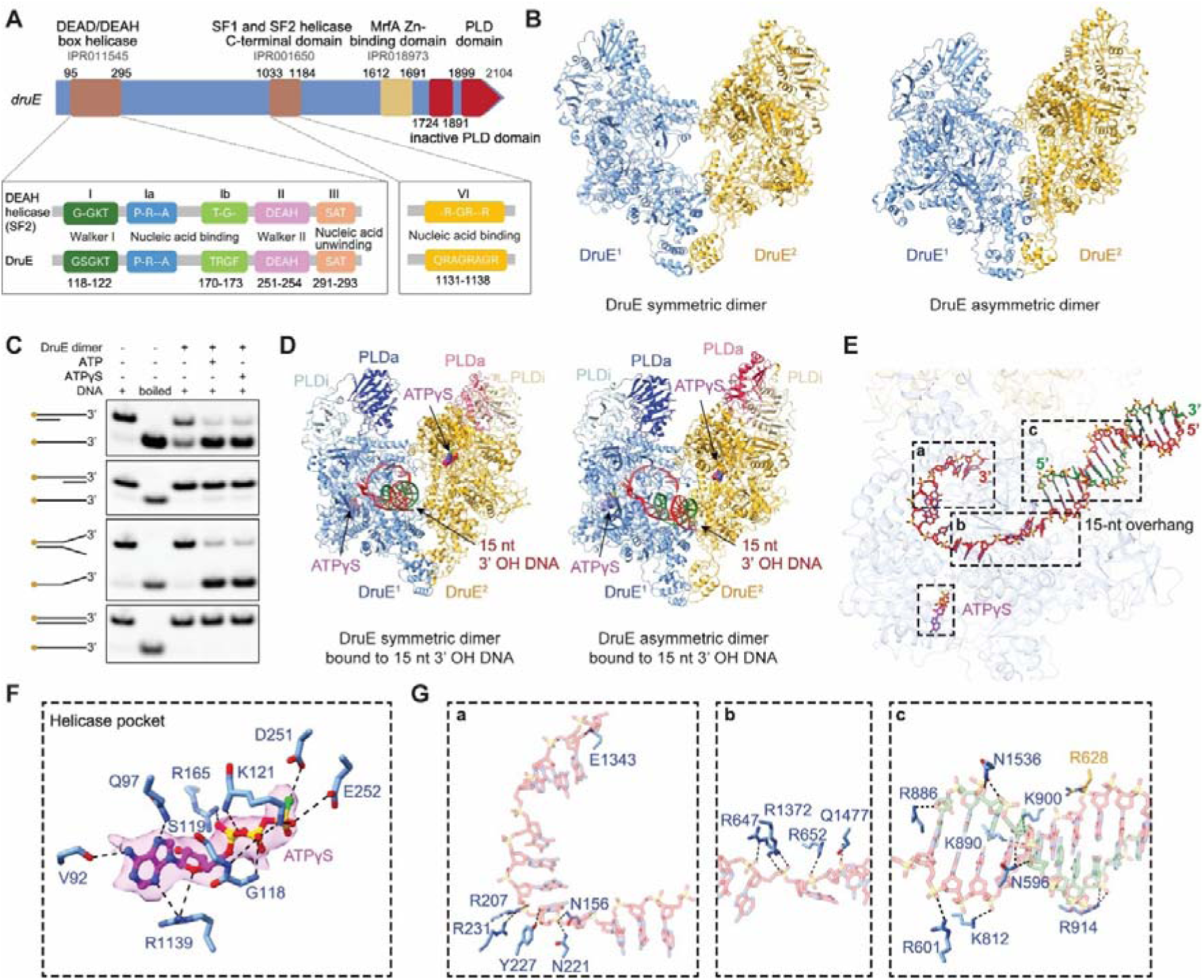
DruE is a dimeric SF2 helicase that preferentially engages 3’-tailed DNA. **(A)** Domain architecture of DruE, showing the SF2 helicase core, SF1/SF2 helicase C-terminal domain, MrfA-like Zn-binding modules, and two C-terminal PLD domains, the upstream inactive and the downstream active one. Insets show the conserved helicase motifs in DruE and their correspondence to canonical SF2 motifs. **(B)** Cryo-EM structures of apo DruE showing two dimeric conformations, a symmetric dimer (left) and an asymmetric dimer (right). The two protomers are coloured blue and gold. **(C)** DNA unwinding assays of DruE using Cy3-labelled DNA substrates. Activity was tested in the presence of ATP or ATPγS, and boiled substrate is shown as a migration control. **(D)** Cryo-EM structures of the symmetric and asymmetric DruE dimers bound to a 15-bp duplex DNA containing a 15-nt 3’ overhang in the presence of ATPγS. The C-terminal PLD domains predicted to be inactive (PLDi) or active (PLDa) are indicated. **(E)** DNA trajectory through the DruE dimer. Dashed boxes highlight three DNA-interaction regions (*a*-*c*) along the substrate path and the ATPγS-binding helicase pocket. **(F)** Close-up view of the helicase nucleotide-binding pocket, showing ATPγS coordinated by surrounding residues. **(G)** Close-up views of the three DNA-interaction regions highlighted in (E), showing residues that contact and stabilise the DNA backbone within the continuous electropositive channel. Region *a* corresponds to the distal single-stranded DNA path, region *b* to the transition from duplex DNA to the 3′ single-stranded overhang, and region *c* to the duplex DNA entry site.

To characterise DruE biochemically, we purified recombinant DruE by Strep-Tactin affinity chromatography followed by size-exclusion chromatography, yielding a predominant species consistent with a stable dimeric assembly (**Figure S3A**). Cryo-electron microscopy (cryo-EM) revealed that DruE adopted a dimeric architecture that exists in two conformational states, corresponding to symmetric and asymmetric dimers (**Figure 3B**, **Table S3**). Despite differences in packing, both assemblies preserved a common overall architecture, indicating that DruE forms a flexible but stable dimeric unit (**Figure S3B-E**). Comparison of these states revealed substantial conformational plasticity within the dimer and between dimers, driven primarily by movements of the Rossmann-like fold within the C-terminal domain (CTD) and adjacent αα-hairpin motifs, resulting in rotational displacement of approximately 39° and positional shifts of ∼42 Å between protomers (**Figure S3F**). These rearrangements indicate that the DruE dimer can adopt multiple conformational states, consistent with the dynamic behaviour typical of SF2 helicases^37,38^.

To determine which DNA substrates were recognised by DruE, we performed DNA unwinding assays using Cy3-labelled substrates designed to probe helicase loading polarity (**Figure 3C**). DruE efficiently unwound 3’-overhang and forked DNA substrates, whereas blunt duplex DNA and 5’-overhang substrates remained largely resistant to unwinding. We observed unwinding in the presence of both ATP and the slowly hydrolysable analogue ATPγS, but not in the absence of nucleotide, indicating that ATP-binding, rather than efficient hydrolysis, was sufficient for productive engagement with these substrates under our assay conditions.

To explore how DruE recognises these substrates, we determined cryo-EM structures of DruE bound to a 15-bp duplex DNA containing a 15-nt 3′ overhang, a substrate that is efficiently unwound in biochemical assays (**Figure 3D**, **Table S3**). In the DNA-bound state, both symmetric and asymmetric dimers were again observed, indicating that DNA binding does not restrict DruE to a single oligomeric configuration. The structures also revealed two ATPγS molecules bound within the canonical helicase nucleotide-binding pockets, one in each protomer (**Figure 3D**). Detailed inspection showed that ATPγS occupies the catalytic cleft formed between the two RecA-like domains of the helicase (**Figure 3E,F**). Residues K121, R165, G118, and S119 contribute to the coordination of the phosphate moiety, whereas V92, K97, and R1139 help define the geometry of the nucleotide-binding pocket. The conserved acidic residues D251 and E252 are positioned appropriately to organise the catalytic environment. Together, these interactions define a compact nucleotide-binding site consistent with the conserved SF2 helicase core.

The DNA substrate traverses a continuous electropositive channel formed primarily within one protomer of the DruE dimer (**Figure 3E**). Along this path, three main interaction regions stabilise the DNA through backbone contacts, consistent with recognition of the DNA architecture rather than the sequence (**Figure 3G**). At the entry site, residues R601, K812, K890, K900, R914, and N1536 contact the duplex DNA backbone, whereas R886 and N596 further stabilise the substrate by interacting with sugar and base moieties. The side chain of R628, contributed by the neighbouring protomer, helps secure the duplex at the channel entrance. The second interaction region corresponds to the transition from duplex DNA to the 3’ single-stranded overhang, where residues R647, R652, and R1372 form a positively charged surface that tracks the phosphate backbone. Q1477 further stabilises DNA orientation. The third region lies near the turning point of the single-stranded overhang, where residues N156, R207, N221, Y227, and R231 contact the DNA, while E1343 stabilises the distal 3’ end (**Figure 3G, panel c**). The position of E1343 also suggests a potential extension of this interaction toward the adjacent DruE protomer within the DruE dimer. In these DNA-bound dimers, the two C-terminal PLD domains, PLDi and PLDa, occupy peripheral positions relative to the helicase core and the main DNA-binding channel (**Figure 3D**), indicating that DNA loading and initial engagement occur through the helicase, whereas the PLD domains remained positioned separately in the dimeric state.

The DNA-bound structures showed that the conserved SF2 helicase motifs (I, Ia, Ib, II, III, and VI; **Figure 3A**) cluster around the nucleotide-binding pocket and the DNA substrate, defining the catalytic helicase core of DruE (**Figure S4A**). In addition, four zinc-binding clusters coordinated by pairs of cysteine residues within the MrfA-like Zn-binding module stabilise structural elements adjacent to the DNA-binding channel (**Figure S4B,C**), suggesting a role in shaping the DNA-engaged state. The homologous DUF1998 module in the DISARM protein DrmB is positioned next to the DrmA helicase core and has been proposed to modulate DrmA activity, suggesting that this Zn-binding module plays a conserved role in tuning defence-helicase function^39^.

Comparison of the DNA-bound and apo DruE structures revealed conformational rearrangements associated with substrate engagement (**Figure S4D-F**). These rearrangements include repositioning of helices, loops and β-motifs that collectively reshape the DNA-binding groove, together with movements in the C-terminal region containing the PLD modules. Additional changes occur in the nucleotide-binding pocket, where residues V92, R165, D251, E252, R1135, and R1139 move toward ATPγS, tightening the catalytic cleft (**Figure S4H,I**). This shift is accompanied by the inward movement of α-helix 156-172 toward ATPγS and displacement of the loop spanning residues 1101-1116 (**Figure S4G,H**), relieving steric clashes present in the apo structure.

Taken together, these findings identify DruE as a dimeric helicase of the YprA-like family that preferentially loads onto exposed 3’ single-stranded DNA and engages its substrate through an extended electropositive channel. The ability to unwind short substrates while tolerating ATPγS further supports a model in which nucleotide binding stabilises the DNA-bound state of DruE, enabling engagement and local strand separation at 3’ ssDNA-containing structures without requiring rapid ATP hydrolysis. The positioning of the PLD domains peripheral to the helicase core further suggests that the nuclease output is not coupled to initial DNA loading in the dimeric state but may instead depend on additional rearrangements or higher-order assembly. Notably, the path of the ssDNA suggests that longer single-stranded regions could extend beyond the primary binding channel and engage the adjacent protomer, suggesting that extended DNA substrates might promote higher-order assembly states of DruE.

### Extended ssDNA can span DruE protomers and promote CTD-mediated higher-order assembly

Because the structure of the DruE dimer complexed with DNA suggested that the ssDNA could extend towards the associated protomer, we investigated how DruE accommodates extended DNA substrates. To this end, we determined cryo-EM structures of DruE bound to a 15-bp duplex DNA containing a longer 28-nt 3’ single-stranded overhang (**Figure 4A**, **Table S3**). DruE again formed both symmetric and asymmetric dimers, with the variability in the effective overhang length reflecting structural heterogeneity (**Figure 4A,B**, **Figure S5A,B**). In the symmetric dimer, the longer substrate was accommodated within the same electropositive channel observed in the 15-nt overhang complex, with continuous density for both the duplex region and the extended single-stranded tail.

**Figure 4.**
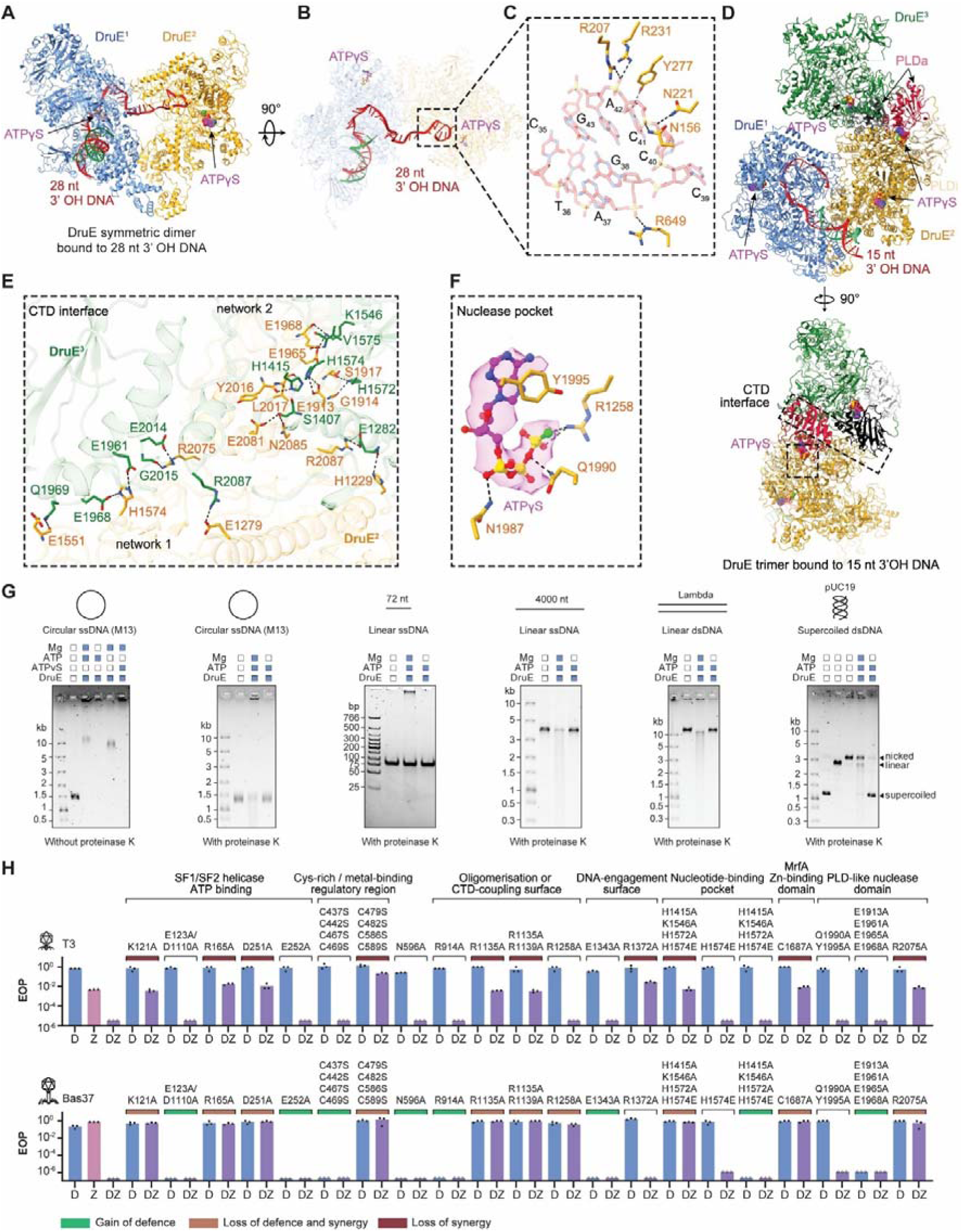
Extended ssDNA promotes higher-order DruE assembly and nuclease activity. **(A)** Cryo-EM structure of the symmetric DruE dimer bound to a 15-bp duplex DNA containing a 28-nt 3′ overhang in the presence of ATPγS. The two protomers are coloured blue and gold. **(B)** Two views of the DNA trajectory through the DruE dimer, showing extension of the 3’ single-stranded tail from one protomer toward the adjacent protomer. The dashed box highlights the region enlarged in (C). **(C)** Close-up view of the extended 3’ ssDNA path, showing residues that contact the DNA, including contributions from the neighbouring protomer. **(D)** Cryo-EM structure of a trimeric DruE assembly bound to a 15-bp duplex DNA containing a 15-nt 3’ overhang. A third protomer (green) associates with the DNA-bound dimer through C-terminal domain (CTD) interfaces, repositioning the C-terminal PLD-containing region within the higher-order assembly. Four ATPγS molecules are observed, two in the helicase pockets and two in CTD-associated pockets, including sites within the active PLD region. **(E)** Close-up view of the CTD interface shown in (D), highlighting the interaction networks that stabilise the trimeric assembly. **(F)** Close-up view of the CTD nuclease pocket, showing ATPγS coordinated by surrounding residues. **(G)** DNA binding and cleavage assays of DruE on the indicated substrates. Left, binding and processing of circular M13 ssDNA in the absence or presence of proteinase K. Right, cleavage assays on 72-nt linear ssDNA, 4000-nt linear ssDNA, linear λ DNA, and supercoiled pUC19 in the presence of Mg²⁺ and ATP, with proteinase K added before analysis. DNA substrates and proteins were used at a final concentration of 5 nM and 250 nM respectively. Where indicated, reactions were supplemented with 5 mM MgCl_2_ and 1 mM ATP or ATPγS. Cleavage products were resolved on 0.7% agarose gels, except for linear ssDNA products, which were analysed on 4-20% TBE polyacrylamide gels. **(H)** Efficiency of plating (EOP) assays for cells expressing Druantia III (D), Zorya II (Z), or both systems together (DZ) carrying the indicated DruE mutants, tested against T3 and Bas37. Mutants are grouped according to the affected structural region or inferred functional class. Colours indicate gain of defence, loss of defence and synergy, or loss of synergy.

Structural modelling indicated that the 28-nt 3’ overhang extends from one protomer towards the neighbouring protomer across the dimer interface (**Figure 4B**). The duplex DNA binds within the basic channel of one protomer, whereas the extended 3′ tail traverses towards the adjacent protomer.

Residues that define the turning point of the ssDNA path in the 15-nt overhang structure (N156, R207, N221, R231, and Y227; **Figure 3G**) also stabilised the extended DNA in the 28-nt complex, together with contacts from the neighbouring protomer (R649) (**Figure 4C**). Thus, while the structure of the DruE complex with the shorter substrate revealed a local bending point, the one with the longer substrate showed that DruE can support a continuous ssDNA trajectory spanning the protomers. These observations indicate that DruE is adapted not only to load onto a 3’ single-stranded region, but also to accommodate extended ssDNA beyond the helicase channel.

Unexpectedly, cryo-EM analysis also revealed a trimeric DruE assembly (**Figure 4D**, **Figure S5C**). In this structure, a third protomer associates with the DNA-bound dimer primarily through CTD-mediated interactions, bringing the C-terminal region containing the PLD domains into an expanded higher-order interface with the protomer not bound to DNA (**Figure 4D**). We also detected higher-order assemblies, including tetramers, although these could not be resolved at high resolution (**Figure S5D**).

Structural comparison indicated that trimer formation requires substantial repositioning of the CTD (∼24.4 Å), relieving steric clashes and enabling recruitment of the third protomer (**Figure S5E,F**). The CTD-CTD interface is stabilised by complementary electrostatic interactions, including insertion of H1574 into an acidic pocket (E1961, E1968, network 1) and reciprocal positioning of E1913 into a histidine-rich pocket (H1415, H1572, H1574, network 2), with additional residues (E1551, E1279, E1965, E1968, E2081, N2085, R2087, and H1229) further reinforcing the interface (**Figure 4E**). These observations indicate that higher-order assembly is accompanied by major rearrangements in the CTD region that contains the PLD modules.

The trimeric structure of DruE contained four ATPγS molecules (**Figure 4D**). Two of these occupied the canonical helicase nucleotide-binding pockets, whereas two additional molecules were located within distinct CTD pockets, defining a previously unrecognised nucleotide-binding site (**Figure 4F**). These two extra ATPγS molecules were coordinated primarily by residues within the active PLD domain, including N1987, Q1990, and Y1995, together with R1258 contributed from an adjacent region of the CTD. The positioning of this pocket along the extended DNA trajectory suggests a potential role in regulating nuclease activity within higher-order assemblies. Together, these observations suggest that the PLD domains are not simply appended to the helicase core but are repositioned during oligomerisation in a manner that could couple higher-order assembly to nuclease activation.

### DruE binds and cleaves diverse long DNA molecules

We tested the nuclease activity of DruE on circular and linear ssDNA substrates. DruE bound strongly to M13 circular ssDNA, forming nucleoprotein complexes that migrated more slowly in agarose gels compared to DruE oligomers not associated with DNA, with Mg^2+^ altering the complex structure but not abolishing binding (**Figure 4G**). Following proteinase K treatment, smearing of the M13 substrate was observed in the presence of ATP and Mg^2+^, indicating that DruE can both bind and cleave long ssDNA (**Figure 4G**).

We next compared the nuclease activity of DruE on short and long linear ssDNA. DruE showed no detectable activity on a 72-nt substrate but efficiently degraded a 4000-nt linear ssDNA molecule (**Figure 4G**). DruE was also active with dsDNA substrates (**Figure 4G**). Linear lambda DNA was efficiently processed in the presence of ATP and Mg^2+^, whereas on supercoiled pUC19, DruE primarily introduced nicks (**Figure 4G**). Consistent with this result, ENDO-pore analysis^40^ revealed recurrent cleavage hotspots without strong sequence specificity (**Figure S5G**). Notably, the DNA-bound structures showed that dsDNA remained positioned at the entrance of the helicase channel, whereas only the single stranded region extends into the catalytic core (**Figure 3E,G**, **Figure 4A**). This observation indicates that DruE does not accommodate intact duplex DNA within the active-site channel. Rather, DNA is likely to be presented locally in a single-stranded or distorted conformation, meaning that only DNA ends, nicks, or locally unwound segments can enter the nuclease active site of DruE. In short, blunt DNA molecules, single-stranded or distorted regions are absent or rare and therefore such substrates cannot support productive DruE activity. In longer DNA molecules, single-stranded regions are more common and therefore longer molecules could support multivalent binding and higher-order assembly of DruE.

Taken together, these results indicate that DruE engages DNA initially as a dimer and can process a broad range of long DNA substrates *in vitro*, including circular ssDNA, long linear ssDNA or linear dsDNA, and supercoiled dsDNA. However, the combined structural and biochemical data suggest that productive activity of DruE is favoured on substrates that provide access to extended or transiently exposed ssDNA, which could facilitate higher-order assembly. This behaviour is reminiscent of cooperative ssDNA-binding proteins such as RecA, which bind ssDNA cooperatively and form substrate length-dependent nucleoprotein filaments^41–43^. Although DruE is structurally distinct and initially loads onto 3’ overhangs through its helicase core, the progressive recruitment of additional protomers along ssDNA suggests a similar transition from a loading state to an extended, assembly-dependent active state. These observations might also explain why Druantia III is sensitive to the loss of RecD. In the absence of RecD, RecBC loads RecA constitutively onto the 3’ ssDNA strand in a Chi-independent manner^29^, which is expected to reduce the accessibility of long ssDNA-containing intermediates. If DruE requires access to such intermediates to load efficiently and assemble into a higher-order active state, then, DruE and RecA are likely to compete, at least transiently, for overlapping substrates.

### Distinct DruE functions underlie Druantia III-Zorya II synergy during Bas37 and T3 infection

To identify the structural features of DruE required for it’s function *in vivo*, we mutated residues highlighted by the DNA-bound structures and by the biochemical evidence that DruE preferentially processes long ssDNA substrates. We then tested the effects of these mutations on the protection against the Druantia-targeted phage Bas37, as well as on the synergy with Zorya II against Bas37 and the Zorya-targeted phage T3 (**Figure 4H**).

Most mutations that reduced synergy also impaired individual Druantia activity, indicating that both outputs rely on the same core functionalities (**Figure 4H**). These included the ATP-dependent helicase activity and CTD-mediated higher-order assembly of DruE. Mutations in the helicase ATP-binding pocket, including K121A and D251A, abolished both Druantia-mediated defence and synergy. The same effect was observed for mutations in additional residues near the ATP-binding site, including R165A and R1135A. Similarly, mutations predicted to disrupt CTD-mediated interactions between adjacent protomers of DruE, including R2075A and the combined H1415A/K1546A/H1572A/H1574E mutant, abolished both Druantia-mediated defence and synergy. These results indicate that productive ATP-dependent remodeling and higher-order assembly are central to DruE function.

The mutational analysis also supported the importance of DNA engagement, but showed that not all DNA-contacting residues contribute equally in all contexts (**Figure 4H**). Mutations in residues positioned near the duplex region, such as N596A and R914A, had little effect, consistent with the structural observation that the duplex is held near the channel entrance whereas the single-stranded region extends into the catalytic path. By contrast, mutation of the ssDNA-contacting residue R1372 strongly impaired individual Druantia activity and synergy against T3, indicating that correct positioning or stabilisation of ssDNA in DruE is important for these outputs. However, R1372A was tolerated during synergy against Bas37, revealing that the requirement for this ssDNA-contacting surface is context-dependent rather than universal. This suggests that proper ssDNA engagement by DruE after initial loading is essential for Druantia alone and for synergy against T3, but not for synergy against Bas37.

Additional mutations further supported this phage-specific separation of DruE functions. R1258A, which maps to the catalytic pocket of the active PLD domain and is positioned adjacent to the phosphate groups within this site, abolished Druantia-mediated defence and impaired synergy against Bas37 but did not affect synergy against T3. These results indicate that DNA cleavage by DruE is required for Druantia-mediated defence and for Bas37-associated synergy, but can be bypassed in the case of synergetic defence against T3, where Zorya likely provides the dominant effector activity.

A subset of mutations enhanced individual Druantia activity, including E1343A and neutralisation of an acidic CTD patch (E1913A/E1961A/E1965A/E1968A), indicating that DruE activity is tuneable (**Figure 4H**). These residues lie along the DNA path or at CTD interfaces and likely influence DNA geometry or assembly stability rather than being directly involved in the activity.

Together, these observations indicate that the functional relationship between Druantia and Zorya depends on the infecting phage. In the case of Bas37 infection, Druantia acts as the primary defence, so that DruE must retain the nuclease activity, as reflected by the requirement for R1258 but not R1372 for synergy. By contrast, in the case of T3 infection, Druantia alone is inactive and synergy reflects a distinct functional relationship between the two defence systems. In this case, DruE appears to contribute primarily through productive engagement of ssDNA-containing DNA intermediates, whereas the dominant effector activity is likely provided by Zorya. Together with the sensitivity of Zorya to factors affecting ssDNA accessibility, these observations are consistent with DruE promoting or stabilising ssDNA structures that are permissive for Zorya activity. Thus, Druantia-Zorya synergy engages DruE in different manners for protection against the two phages tested in this work, with Bas37-associated synergy retaining the requirement for the DruE effector output, whereas T3-associated synergy relies primarily on DruE-mediated DNA engagement.

### DruH is a likely phage sensor with dimerization-dependent nuclease activity

We next investigated the role of DruH, the second component of Druantia III that is specific to this Druantia subtype^13^. Sequence analysis assigned DruH to the YjiT family (**Figure 5A**), a poorly characterised protein family whose *E. coli* representative is encoded within the immigration control region, a highly variable chromosomal locus enriched in restriction functions^44^. This genomic context is consistent with a role in defence against incoming DNA.

**Figure 5.**
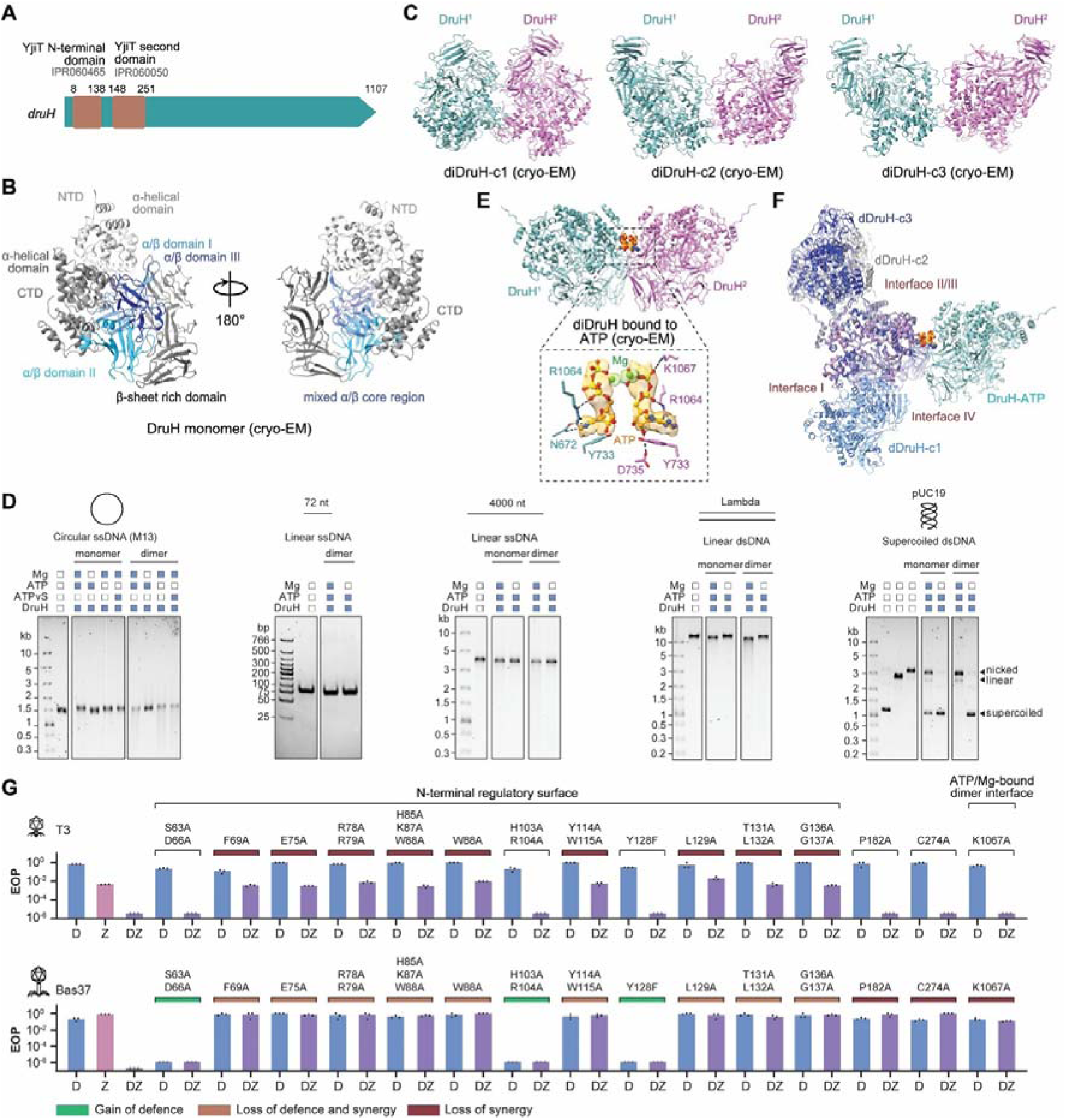
DruH is a phage-responsive factor whose nuclease activity is enhanced by dimer formation. **(A)** Domain architecture of DruH, showing the predicted YjiT N-terminal and second domains. **(B)** Cryo-EM structure of the DruH monomer shown in two views. Major structural regions are indicated, including the N-terminal domain (NTD), C-terminal domain (CTD), α-helical domain, β-sheet-rich domain, and the mixed α/β core containing α/β domains I-III. **(C)** Cryo-EM structures of three DruH dimer conformations, diDruH-c1, diDruH-c2, and diDruH-c3. The two protomers are coloured cyan and magenta. **(D)** Nuclease assays of monomeric and dimeric DruH on the indicated DNA substrates in the presence of Mg^2+^ and ATP or ATPγS, as shown. **(E)** Cryo-EM structure of the DruH dimer bound to ATP and Mg^2+^. The two protomers are coloured cyan and magenta. The inset shows the composite nucleotide-binding site at the dimer interface. **(F)** Structural comparison of the ATP-bound dimer with the three apo dimer conformations, highlighting four alternative dimer interfaces (interfaces I-IV) observed across the DruH assemblies. **(G)** Efficiency of plating (EOP) assays for cells expressing Druantia III (D), Zorya II (Z), or both systems together (DZ) carrying the indicated DruH mutants, tested against T3 and Bas37. Mutants are grouped according to the affected structural region or inferred functional class. Colours indicate gain of defence, loss of defence and synergy, or loss of synergy.

To gain insight into DruH function, we compared its structure to known protein folds. The N-terminal region of DruH (residues 1-676) showed structural similarity to phage baseplate wedge proteins of the J family, with structural superposition revealing a shared core with an RMSD of 1.15 Å over 171 pruned Cα atom pairs (**Figure S6A**). Within this region, DruH contains three immunoglobulin (Ig)-like domains (residues 257-464, 465-631, and 632-748) (**Figure S6B**). Because Ig-like domains typically mediate protein-protein interactions, in particular, those involved in molecular recognition^45^, whereas J-like baseplate proteins function as interaction scaffolds within the phage baseplate^46,47^, these combined structural features suggest that DruH functions as a protein-recognising infection sensor. This possibility is consistent with the results of co-immunoprecipitation assays, which detected a weak association between DruH and the phage baseplate component Bas37_0273 (**Figure S2I**). Notably, Bas37_0273 is a homolog of T4 gp7, which interacts with the J-like T4 gp6 protein within the phage baseplate^47,48^.

Several additional observations support this interpretation. Mass photometry analysis of DruH and DruE did not reveal stable interactions between the two proteins (**Figure S6C**), arguing against a constitutive DruE-DruH complex. Moreover, DruH proteins were substantially more divergent than their cognate DruE partners (**Figure S6B,D**), suggesting a stronger phage-driven positive selective. In two independent examples where a single DruE variant co-occurred with multiple, albeit closely related DruH variants, the FEL^49^ and FUBAR^50^ methods implemented in HyPhy^51^ identified multiple residues in DruH inferred to evolve under positive selection, several of which mapped to the Ig-like domains (**Figure S6E**). Together, these findings strongly suggest that DruH is the phage-sensing component of Druantia III.

We next examined the structural and biochemical properties of DruH. Recombinant DruH resolved into monomeric, dimeric and aggregated species by size-exclusion chromatography (**Figure S6F**). Cryo-EM revealed a monomeric DruH structure together with three distinct apo dimeric conformations (diDruH-c1-3) (**Figure 5B,C**, **Figure S6G**, **Table S3**). The monomer adopts a multidomain architecture in which N-terminal and C-terminal α-helical regions flank a central core of three α/β domains (I-III), and a β-sheet-rich region adjacent to the CTD. In the dimer configurations, the two DruH protomers associate through distinct interfaces, indicating substantial conformational plasticity within the assembly (**Figure 5C**, **Figure S6G**).

Given the association of YjiT-family proteins with defence against invading DNA, we speculated that the large C-terminal region of DruH distinct from the J-like/Ig-like scaffold implicated in infection sensing, might contribute an additional function, perhaps, being directly involved in DNA-processing during defence. Monomeric DruH showed only weak nuclease activity with all the substrates tested, whereas dimeric DruH was markedly more active, particularly on dsDNA substrates, with the clearest effect with the supercoiled pUC19 (**Figure 5D**). In reactions with monomeric DruH, a substantial fraction of the supercoiled substrate remained intact, whereas dimeric DruH cleaved all the substrate, yielding, predominantly, nicked products, with a smaller amount of linear DNA. These results indicate that the nuclease activity of DruH is strongly enhanced in the dimeric state.

To capture a DNA-bound state, we incubated DruH with supercoiled pUC19, but although cryo-EM revealed particles associated with DNA filaments **(Figure S6H**), no defined DNA-bound structure was resolved. Instead, we identified a distinct DruH dimer bound to ATP and Mg^2+^ that was structurally distinct from the apo assemblies (**Figure 5E**, **Table S3**). In this structure, two ATP molecules lie at the dimer interface and are coordinated by two Mg^2+^ ions. The nucleotide-binding site is assembled across the dimer interface. In each protomer, R1064 from the CTD of one protomer and R1064/K1067 from the opposing CTD are positioned to contact the phosphate groups of the bound ATP molecules, whereas N672, Y733 and D735 provide additional contacts to the nucleotide-binding pocket, consistent with stabilisation of ATP binding (**Figure 5E**). The presence of this composite nucleotide-binding site indicates that ATP/Mg^2+^ binding is associated with a distinct dimeric state.

To further examine how DruH might interact with DNA, we generated AF3 models of dimeric DruH bound to dsDNA and ssDNA (**Figure S6I,J**). In both models, the nucleic acid tracks along a surface formed primarily by the N-terminal α-helical regions of the two protomers. The predicted contacts are contributed by both protomers, consistent with a cooperative mode of substrate engagement by the dimer (**Figure S6K**). Structural comparison of these models with the cryo-EM structure of the ATP/Mg^2+^-bound dimer suggested substantial repositioning of the N-terminal regions, consistent with the conformational plasticity observed the experimentally resolved structures of DruH dimers (**Figure S6L**).

Comparison of the structures of the DruH dimers identified four alternative intermolecular interfaces (interfaces I-IV, **Figure 5F**, **Figure S6M**), consistent with substantial conformational flexibility. Although the present data do not establish which dimer structures is predominant during defence, they support a model in which DruH samples multiple assembly states, and nucleotide binding is associated with a distinct dimeric configuration linked to DNA-processing.

Together, these findings suggest that DruH combines a phage-sensing module with a dimerization-dependent nuclease activity, in which phage recognition by DruH could promote or stabilise a distinct dimeric state associated with nucleotide binding and nuclease activity.

### A regulatory surface of DruH controls pathway activation

To identify functionally important regions of DruH, we introduced mutations into residues highlighted by structural and evolutionary analyses and tested their effects on Druantia-mediated defence and on the synergy with Zorya (**Figure 5G**). The mutations segregated into three functional classes.

The largest class of substitutions (F69A, E75A, R78A/R79A, H85A/K87A/W88A, W88A, Y114A/W115A, L129A, T131A/L132A, G136A/G137A) abolished both standalone Druantia activity and synergy, identifying an N-terminal surface in DruH spanning, approximately, residues 63-137 as a key functional region required for the pathway output. Within the same region, a second class of substitutions (S63A/D66A, H103A/R104A, Y128F) enhanced Druantia-mediated defence against Bas37. The co-existence of loss- and gain-of-function within the same N-terminal surface indicates that this region does not simply provide a constitutive structural or binding function but also modulates activation of DruH, consistent its role in the early steps of the defence pathway, upstream of DNA processing.

The third class of mutations (P182A, C274A, K1067A) preserved the individual defence activity of Druantia III alone but selectively impaired synergy with Zorya II against Bas37, while having little effect on the synergy against T3. P182A and C274A are located outside the main N-terminal functional patch, whereas K1067 maps to the nucleotide-coupled dimer interface identified in the ATP/Mg^2+^-bound structure. Because T3 escapers remain sensitive to Zorya II alone while escaping the additional protection conferred by the combined systems (see above), both phages most likely engage the Druantia pathway. Thus, the different effects of these mutations on Bas37- and T3-associated synergy indicate that DruH is subject to distinct functional requirements in the two contexts. At present, we cannot distinguish whether this reflects differences in trigger recognition, in the activated states adopted by DruH, or in downstream coupling to DNA-processing activity.

Together, these results define an N-terminal regulatory surface in DruH that is required for pathway activation and can either promote or suppress the defence output, as well as a nucleotide-coupled dimer interface that contributes to defence in a phage-dependent manner. These findings suggest that DruH acts as an upstream regulatory component of the Druantia pathway, while also indicating that the mechanistic requirements for DruH are phage-specific.

### Synergistic defences against Bas37 and T3 require distinct ZorE states

To further investigate the synergy between Druantia III and Zorya II, we tested whether ZorE forms a stable complex with the Druantia proteins. We could not detect stable interactions under the conditions tested (**Figure S7A**). Although not excluding transient, weak, or trigger-dependent contacts *in vivo*, these observations argue against a constitutive complex between the two systems and instead suggest that synergy arises from functional coupling of their activities. We therefore asked which structural features of ZorE are required for standalone Zorya defence and for synergy with Druantia.

Sequence analysis identified an N-terminal DUF8245 domain and a central HNHc nuclease domain in ZorE (**Figure 6A**). As described above, the subset of ZorE proteins associated with Druantia III also frequently carries a C-terminal nuclease-like extension (**Figure S1D**), suggesting that this domain contributes to the role of ZorE in the synergy with Druantia III.

**Figure 6.**
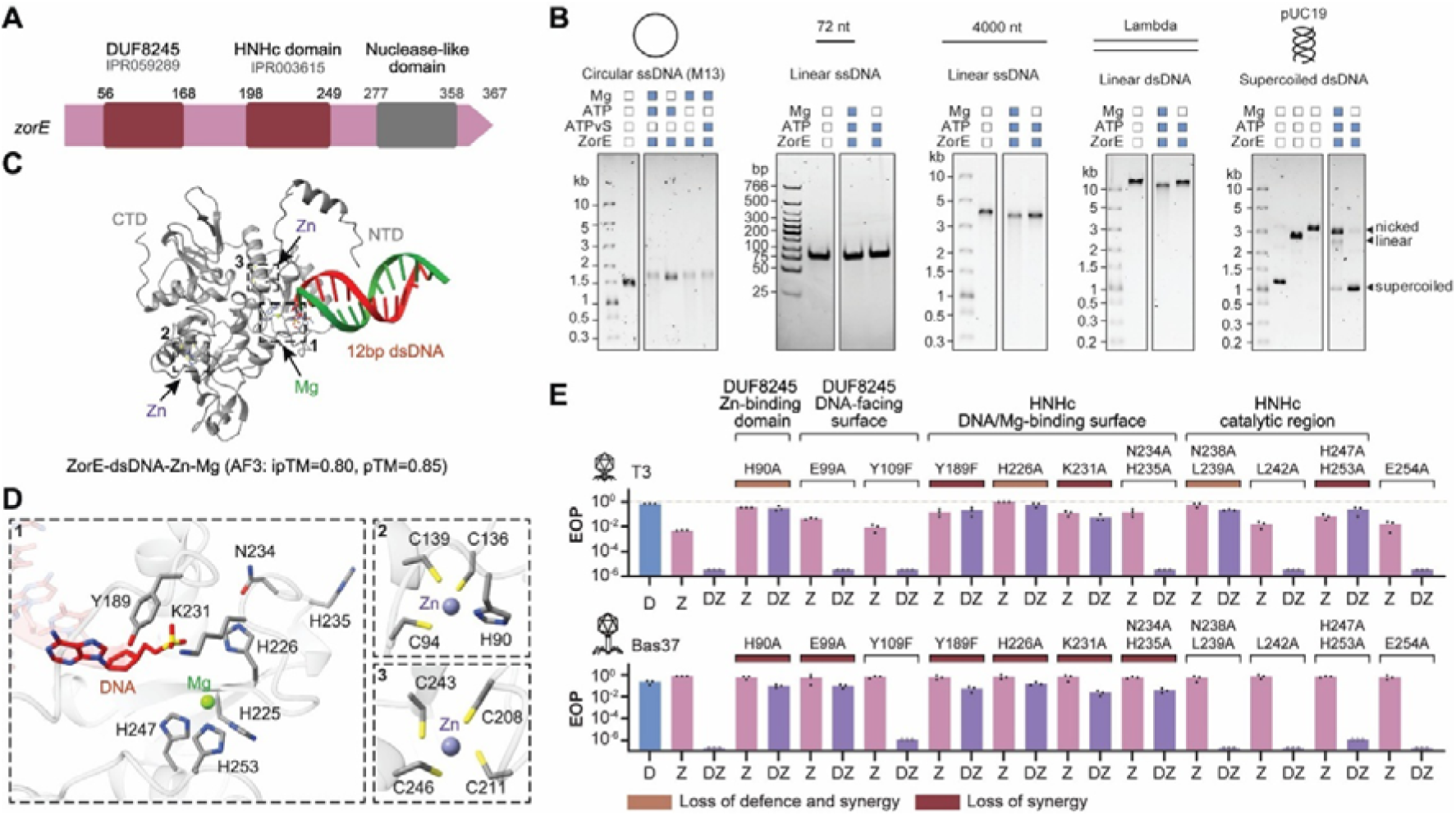
Shared and phage-specific requirements for ZorE activity in Zorya II defence and Druantia III-Zorya II synergy. **(A)** Domain architecture of ZorE, showing the DUF8245 domain, HNHc domain, and C-terminal nuclease-like domain. **(B)** Nuclease assays of ZorE on the indicated DNA substrates. DNA substrates and ZorE were used at final concentrations of 5 nM and 250 nM, respectively. Where indicated, reactions were supplemented with 5 mM MgCl_2_ and 1 mM ATP or ATPγS. Reaction products were resolved on 0.7% agarose gels, except for linear ssDNA products, which were analysed on 4-20% TBE polyacrylamide gels. **(C)** AlphaFold3 (AF3) model of ZorE bound to a 12-bp dsDNA substrate in the presence of Mg^2+^ and Zn^2+^. The N-terminal domain (NTD), C-terminal domain (CTD), DNA, and metal ions are indicated. **(D)** Close-up views of the regions highlighted in (C), showing the predicted DNA/Mg-binding region (1) and the two Zn-binding sites (2, 3). **(E)** Efficiency of plating (EOP) assays for cells expressing Druantia III (D), Zorya II (Z), or both systems together (DZ) carrying the indicated ZorE mutants, tested against T3 and Bas37. Mutants are grouped according to the affected structural region or inferred functional class. Colours indicate loss of defence and synergy or loss of synergy.

To guide mutagenesis, we generated an AF3 model of ZorE bound to dsDNA, given that ZorE has been previously shown to possess a nicking activity^15^, which we confirmed here with a supercoiled dsDNA substrate (**Figure 6B**). We also observed activity against circular ssDNA although the activity with supercoiled dsDNA was higher (**Figure 6B**). Together with the DinI and RecD phenotypes described above, these findings suggest that ZorE cleaves substrates that combine dsDNA with local ssDNA exposure or topological stress. In the AF3 model, the N-terminal and C-terminal regions flank the DNA duplex, and the predicted DNA-interacting surface is supported by two Zn-binding sites together with a Mg^2+^ ion positioned adjacent to the substrate (**Figure 6C,D**). Residues Y189, H226, K231, N234 and H235 cluster along the predicted DNA-interacting surface adjacent to the Mg^2+^ ion, whereas H247 and H253 flank the neighbouring metal-associated region of the HNHc domain. The model also places H90 within an N-terminal Zn-binding site and positions E99 and Y109 close to the predicted DNA-facing surface of the DUF8245 region. Although these assignments remain predictive, they provided a useful framework for testing how distinct regions of ZorE contribute to standalone Zorya defence and to synergy with Druantia III against different phage classes.

The first class of mutations, including H90A, Y189F, H226A and K231A, abolished both standalone Zorya defence and synergy against all phages tested (**Figure 6E**). These residues span both the N-terminal DUF8245 region and the central HNHc domain, indicating that both are required for the individual antiviral activity of Zorya and for its contribution to synergy. H226 is a likely catalytic residue because histidines at equivalent positions in HNH nucleases commonly belong to the catalytic centre, whereas lysines, such as K231, are thought to stabilise the DNA-engaged active conformation and/or help position the scissile phosphate^52^. The strong phenotypic effect of Y189F further suggests that the hydroxyl-bearing side chain at this HNH-adjacent position is functionally important, consistent with a role in local hydrogen-bonding and/or coupling between the flanking region and the nuclease core^53^. H90, which maps to the predicted Zn-binding site in the N-terminal region, likely contributes to the structural integrity of the broader DNA-engaged state required for both individual defence and synergy.

The second class of mutants revealed selective requirements for the synergy against Bas37 (**Figure 6E**). E99A and N234A/H235A abolished synergy against Bas37, but not against T3, although both mutations modestly reduced standalone Zorya activity against T3. E99 lies within the N-terminal DUF8245 domain, whereas N234/H235 are within the HNHc domain near the predicted DNA/Mg^2+^-interacting surface. In other HNH nucleases, similarly located residues often help organise the metal-bound active-site geometry^54^. The phenotype of N234A/H235A therefore suggests that Bas37-associated synergy depends on a particular DNA-engaged or catalytically poised conformation of the HNHc domain of ZorE. The effect of E99A further indicates that synergy against Bas37 also depends on contributions from the N-terminal region of ZorE.

The third class of mutants showed the opposite behaviour (**Figure 6E**). N238A/L239A and H247A/H253A in the HNHc domain abolished standalone Zorya defence as well as synergy against T3 but did not impair synergy against Bas37. In the AF3 model, H247 and H253 flank the predicted Mg^2+^-binding site, whereas N238/L239 map to the adjacent HNHc surface. These phenotypes indicate that defence against T3 requires the cleavage-competent conformation of the HNHc domain, whereas synergy against Bas37 tolerates disruption of features that are critical for the standalone defence activity of Zorya II. A possible explanation of these findings is that ZorE engages distinct, but related DNA states in the synergistic defence against the two phages. Under this scenario, T3 defence depends on the nuclease-competent HNHc conformation, whereas synergy against Bas37 engages ZorE in a different functional state that also depends on the HNHc active site organisation but is less tightly coupled to the canonical catalytic configuration required for the standalone Zorya activity.

Taken together, these results show that ZorE does not contribute to synergistic defences through a single mechanism. In the case of T3, ZorE remains closely coupled to the canonical Zorya pathway, consistent with the requirement for ZorAB (**Figure 1B**) and with the ZorE mutant profile indicating dependence on the intact HNHc configuration required for the standalone Zorya defence. These findings align with the results of the mutational analysis of DruE, which showed that synergy against T3 depended on DruE-mediated engagement of ssDNA-containing intermediates but not on the nuclease activity of DruE (**Figure 4I**). These observations indicate that Zorya provides the dominant effector activity against T3, whereas DruE helps generate, stabilise, or preserve a DNA state permissive for ZorE action.

By contrast, synergy against Bas37 requires ZorE but not ZorAB (**Figure 1B**), indicating that in this case, the synergistic interaction with Druantia III does not depend on direct phage-triggered activation of Zorya II via the canonical route. These observations, again, mirror the DruE mutant phenotypes, which showed that synergy against Bas37 required the nuclease activity of DruE. Thus, whereas synergy against T3 is led by Zorya, employing DruE primarily for DNA engagement, synergy against Bas37 is, conversely, led by Druantia, co-opting ZorE in a distinct functional state. Druantia appears to be the dominant defence arm against Bas37, whereas ZorE likely acts downstream on a related but non-identical DNA substrate state that no longer requires the canonical ZorAB-dependent defence route. More specifically, Druantia might generate or expose a DNA substrate that can be productively targeted by ZorE without direct phage-triggered activation through ZorAB, potentially by creating or preserving structured DNA intermediates that remain accessible in the presence of host DNA-processing factors such as RecA.

Together, these findings indicate that Druantia III-Zorya II synergy does not arise through simple co-activation of two complete defence pathways. Instead, the two cases of this synergy explored here involve phage-specific rewiring of the interactions between the DNA-processing activities of the two systems.

## Discussion

This study defines a mechanistic framework for phage defence by Druantia III and explains how Druantia III and Zorya II synergise through two distinct, phage-specific pathways. Rather than simple co-activation of two complete defence systems, our findings show that the two defence pathways converge on DNA intermediates generated during infection and engage them differently depending on the phage (**Figure 7**).

**Figure 7.**
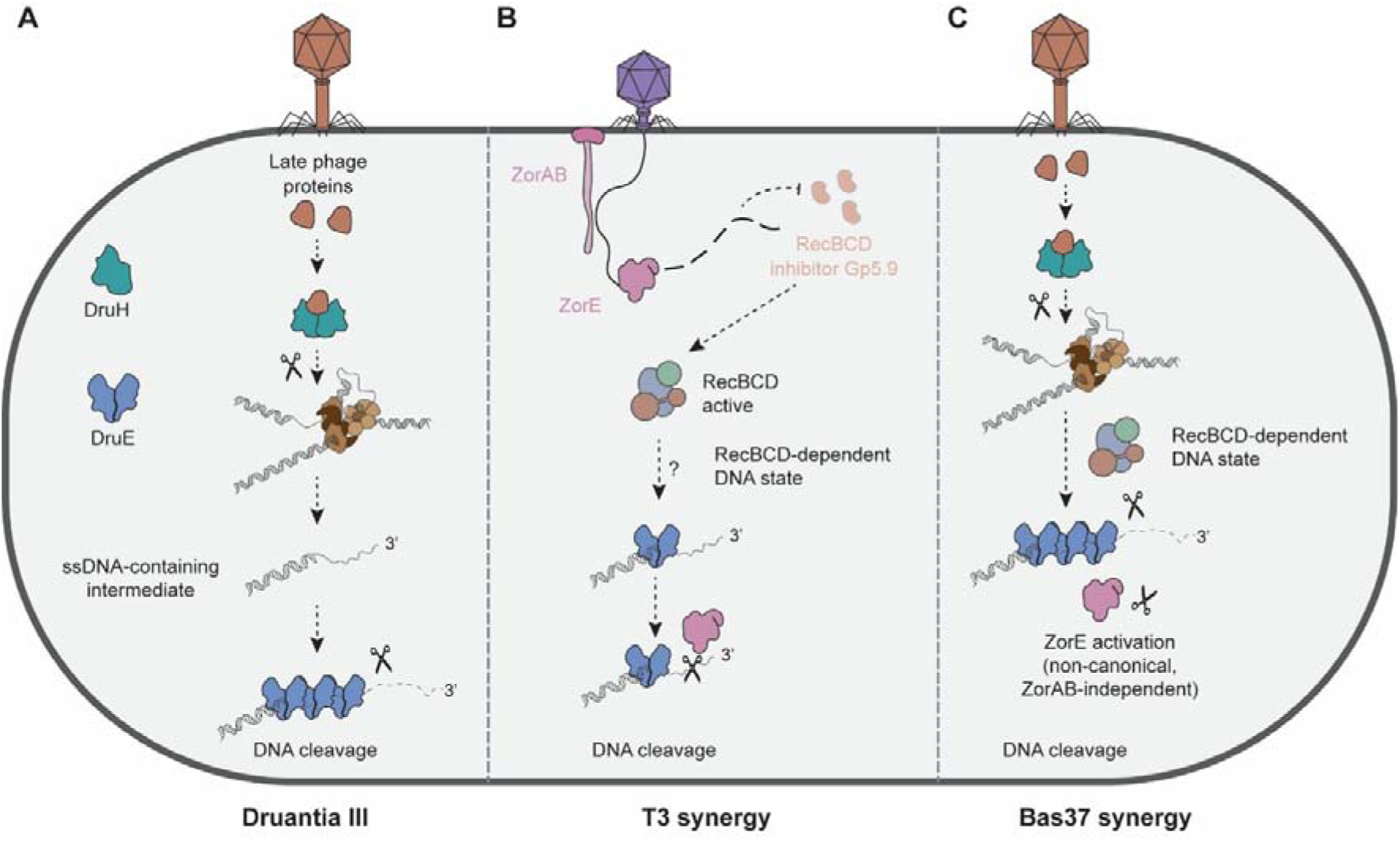
Proposed models for Druantia III activity and phage-specific synergy with Zorya II. **(A)** Model for Druantia III activity. Late phage proteins are proposed to activate DruH during infection. Activated DruH then acts on infection-associated phage DNA structures, likely generated during late stages of replication, leading to the formation or exposure of an ssDNA-containing intermediate. DruE loads onto this intermediate through its helicase core, assembles into a higher-order state on extended ssDNA, and promotes downstream DNA cleavage. **(B)** Model for synergy during T3 infection. During T3 infection, Zorya II is activated through the canonical ZorAB pathway, but the phage-encoded RecBCD inhibitor Gp5.9 impairs the RecBCD-dependent DNA state required for synergy. Zorya is proposed to act early enough to limit the accumulation or action of Gp5.9, thereby preserving RecBCD activity. This allows formation or maintenance of a Druantia III- and RecBCD-dependent DNA state that is permissive for ZorE activity, resulting in synergistic DNA cleavage. **(C)** Model for synergy during Bas37 infection. During Bas37 infection, Druantia III provides the dominant pathway, as described in panel (A). Late phage proteins activate DruH, leading to processing of infection-associated DNA structures and formation of a DruE- and RecBCD-dependent DNA state that supports ZorE recruitment in a non-canonical, ZorAB-independent manner. In this branch, ZorE acts as an additional effector layered onto the Druantia pathway, resulting in enhanced DNA cleavage.

In the case of Druantia III functioning on its own, DruH most likely acts upstream as the phage sensor, whereas DruE functions as the effector, providing the helicase and nuclease activities required for defence. The case for DruH as the sensor remains indirect, but this role is supported by the presence of Ig-like and J-like domains implicated in protein recognition, by the detected weak association with the Bas37 baseplate component, and by the enrichment of positively selected residues in the Ig-like region. Together, these observations are consistent with DruH participating directly in the recognition of phage-encoded trigger proteins. DruE is a YprA-like SF2 helicase-nuclease that loads onto 3’ ssDNA-containing DNA through its helicase core, engages DNA through an extended electropositive channel, and forms higher-order assemblies on extended ssDNA. These properties suggest that DruE first recognises DNA architecture rather than sequence and then transitions, upon oligomerisation, into a nuclease-competent state, in line with the broader view of YprA-family defence helicases as enzymes specialised in recognising and processing infection-associated DNA structures rather than unique sequence motifs^16^.

Taken together, these observations suggest that Druantia III acts on late infection-associated DNA structures. Given that DruH is triggered by late phage structural proteins and dimeric DruH shows the strongest nuclease activity with topologically complex DNA substrates, DruH most likely acts on branched, nicked, or otherwise stressed DNA intermediates generated during late phage replication. This scenario is especially plausible for Bas37, a T4-like phage, because late T4-like replication becomes recombination-dependent and proceeds through branched intermediates^55^. DruH activity would then generate or expose DNA molecules containing accessible ssDNA, onto which DruE could load through its helicase core. Once engaged, DruE can extend across longer ssDNA regions, recruit additional protomers, and transition into a higher-order nuclease-competent assembly. The genetic data further indicate that Druantia III is selectively sensitive to the loss of RecD. In the absence of RecD, RecBC loads RecA constitutively onto 3’ ssDNA in a Chi-independent manner^29^, which is expected to reduce the accessibility of ssDNA-containing intermediates. Thus, the Δ*recD* phenotype is most readily explained by the reduced access to the ssDNA-rich DNA structures on which DruE assembles. Thus, the key substrate for DruE apparently is not a particular DNA sequence, but an infection-associated DNA architecture that combines duplex DNA with exposed ssDNA and is of sufficient length to support assembly (**Figure 7A**). By contrast, the synergistic response requires intact RecBCD, indicating that full RecBCD activity becomes specifically important at the point where Druantia III function is coupled to that of Zorya II.

Our findings also help place Druantia III in the context of the broader Druantia family. DruE is the only component shared across Druantia subtypes, suggesting that DNA processing by a DruE-like helicase-nuclease is the conserved core function of Druantia, whereas the subtype-specific partner proteins determine how that core is activated. In Druantia type III, DruH appears to provide this upstream trigger-recognition and regulatory layer.

Our data further suggest that Zorya II is most efficient when ssDNA-containing intermediates are not occupied by RecA and thus remain accessible. This conclusion is supported by the DinI and Δ*recD* phenotypes, which together point to an inhibitory effect of RecA loading on the activity of Zorya II. It is also consistent with the biochemical properties of ZorE, which acts more efficiently on conformationally complex DNA substrates. RecA binding would inhibit Zorya by coating ssDNA-rich intermediates and reducing access to the structured DNA states on which ZorE acts most productively.

The Druantia III-Zorya II synergy against T3 can be understood within this framework. T3 escaper phenotypes show that Druantia directly responds to T3, indicating that Zorya is not required for Druantia activation in this case. However, synergy is abolished by the loss of any of the RecBCD subunit and by expression of phage RecBCD inhibitors Gam and Gp5.9, indicating that the shared synergistic state requires intact RecBCD. We therefore propose that during T3 infection, Zorya acts early enough^14,15^ to limit the accumulation of Gp5.9 and thereby preserve RecBCD activity. This protective effect of Zorya II likely allows RecBCD to collaborate with Druantia III in generating the DNA state permissive for the ZorE activity and therefore synergy. Mutational data indicate that, in this defence branch, DruE contributes primarily through DNA engagement rather than through its own nuclease activity, whereas Zorya II provides the dominant effector output (**Figure 7B**).

The synergy against Bas37 follows the same overall logic but with the two systems contributing in the opposite order. In this case, Druantia is the system that first senses infection, ZorAB is dispensable, and the main effector activity is provided by Druantia. Our data support a model in which Druantia III acts first on late infection-associated DNA structures, and RecBCD participates at a subsequent DNA-processing step that links Druantia III activity to ZorE recruitment. In this context, DruE nuclease activity remains important, whereas ZorE appears to be co-opted in a distinct functional state that no longer requires activation by ZorAB. Thus, during Bas37 infection, Druantia provides the dominant active pathway, whereas ZorE is recruited secondarily through a related but non-identical DNA structure (**Figure 7C**). These observations further raise the possibility that in the canonical Zorya II defence context, activation of ZorE by ZorAB is linked to the generation of DNA structures optimal for the activity of ZorE.

These two forms of synergy can therefore be unified within a single model centred on a shared RecBCD-dependent DNA-processing hub. In both cases, RecBCD, together with Druantia III, contributes to the formation or maintenance of DNA states permissible to ZorE activity, RecA competes for access to these intermediates, and ZorE acts most efficiently when they are not occupied by RecA. The main difference lies in how the two systems contribute once this shared hub is reached. During T3 infection, Zorya preserves the conditions needed for the hub to remain accessible for defence and provides the key ZorE effector step, whereas Druantia contributes primarily through DNA engagement. By contrast, during Bas37 infection, Druantia shapes the relevant DNA substrate more directly whereas ZorE is engaged secondarily in a non-canonical, ZorAB-independent manner.

Together, these findings define a mechanistic model for Druantia III and show how synergy with Zorya II can emerge through shared DNA intermediates albeit in a phage-specific manner. More generally, this work shows that bacterial defence strategies can be shaped not only by the set of systems present in a cell, but also by how those systems of their individual components access, preserve, or modify shared infection-generated DNA states. These results illustrate a broader principle in which such DNA intermediates can act as integration points for synergistic interactions among defence systems.

## Acknowledgements

We thank Alexander Harms (ETH Zurich) for kindly providing the BASEL collection of phages, Kotaro Kiga (Jichi Medical University) for providing phage T5, and Giuseppina Mariano (University of Glasgow) for kindly providing plasmids. Y.W. was supported by an NIHR Southampton Biomedical Research Centre Postdoctoral Bridging Fellowship. F.N. was supported by a Wessex Health Partners and National Institute for Health and Care Research Wessex Experimental Medicine Network seed fund. S.K.G. and E.V.K. were supported by the Intramural Research Program of the National Library of Medicine, NIH. D.J.P. was supported by NIH grant GM145888, the Maloris Foundation, and the Memorial Sloan Kettering Cancer Center Core Grant (P30-CA008748). In addition to the MSKCC cryo-EM facilities, portions of this work were performed at the National Center for CryoEM Access and Training (NCCAT) and the Simons Electron Microscopy Center at the New York Structural Biology Center, supported by the NIH Common Fund Transformative High Resolution Cryo-Electron Microscopy Program (U24 GM129539), the Simons Foundation (SF349247), and the New York State Assembly. This work utilized the computational resources of the NIH HPC Biowulf cluster (http://hpc.nih.gov). The contributions of the NIH authors are considered Works of the United States Government. The findings and conclusions presented in this paper are those of the authors and do not necessarily reflect the views of the NIH or the U.S. Department of Health and Human Services.

## Author contributions

Conceptualisation, F.L.N.; Methodology; Y.W., Z.Z., S.K.G., P.W., E.V.K., D.J.P., F.L.N.; Formal Analysis, Y.W., Z.Z., S.K.G., P.W., F.L.N.; Investigation, Y.W., Z.Z., S.K.G., P.W.; Visualisation, Y.W., Z.Z., S.K.G., F.L.N.; Writing – Original Draft, Y.W., Z.Z., S.K.G., D.J.P., F.L.N.; Writing – Review & Editing, all authors; Resources, P.W., E.V.K., D.J.P., F.L.N.; Funding Acquisition, E.V.K., D.J.P., F.L.N..

## Methods

### Bacterial strains and phages

*E. coli* strain DH5α was used to clone plasmids pACYCDuet-1, 8A, pCOLADuet-1 or pCDFDuet-1 with individual defence systems and phage genes. *E. coli* BL21-AI cells containing plasmid(s) with the defence systems were used for phage assays. All bacterial strains were grown at 37 °C in Lysogeny Broth (LB) with 180 rpm shaking for liquid cultures, or in LB agar (LBA) plates for solid cultures. Strains containing plasmid pACYCDuet-1, 8A, pCOLADuet-1 or pCDFDuet-1 were grown in media supplemented with 25 µg/ml of chloramphenicol, 100 µg/ml of ampicillin, 50 µg/ml of kanamycin or 20 µg/ml of spectinomycin, respectively. Recombinant proteins were expressed in *E. coli* BL21 (DE3) cells. Cells were cultured in LB supplemented with the appropriate antibiotics at 37 °C with shaking at 220 rpm. Phages used in this study and their origins are described in **Table S4**. All phages were produced in LB with their host strain, centrifuged, filter-sterilized, and stored as phage lysates at 4°C.

### Phylogenetic analysis of Druantia and Zorya

All DruE, DruH, and ZorE proteins were retrieved from precomputed PADLOC^18,19^ annotations of RefSeq (release 209)^17^. For DruE and DruH, we applied filtering criteria requiring an e-value < 1e−10 and coverage of at least 70% of the corresponding PADLOC profile. Additionally, records originating from genomes that had been removed from RefSeq as of April 15, 2025 were excluded. Only proteins belonging to complete defence systems were retained. Non-redundant protein sequences were downloaded from RefSeq and clustered using MMseqs2^56^ easy-cluster with the following parameters: -c 0.9 --min-seq-id 0.98 --cov-mode 0. One representative sequence per cluster was selected for downstream analysis. Multiple sequence alignments of representative sequences were generated using MUSCLE v5^57^ with the --super5 option. For ZorE, alignments were manually inspected, and clusters corresponding to inactive variants were removed (47 clusters in total). The remaining sequences were then realigned. All alignments were trimmed using pytrimAl with the --gappyout option^58^. Trimmed alignments were used for phylogenetic reconstruction with IQ-TREE2 v2.1.3^59^ with -B 1000. The resulting phylogenetic trees were visualised using the ggtree package in R^60^. Taxonomic information for all genomes was retrieved using NCBI Datasets command-line tools^61^ based on species names from the NCBI assembly summary. Mobile genetic elements (MGEs) were identified as follows: prophages and plasmids were predicted using geNomad^62^, and genomic islands were predicted using TreasureIsland^63^. Coordinates of MGEs were intersected with defence system loci using custom R scripts. Initial clade delineation on DruE and DruH phylogenetic trees was performed using TreeCluster^64^ with the -m med_clade method and thresholds of -t 3 for DruE and -t 2.4 for DruH. These clades were manually inspected and merged, guided by the minimal clade structure observed in the DruH phylogeny. Concordance between DruE-DruH and DruE-ZorE phylogenies was visualised using custom R script. Bacterial habitats associated with individual clades in the DruE and DruH phylogenies were inferred using species-level classifications from ^65^. Species in our dataset were matched to records in the SPIRE v1 database^66^. If species-level annotations were unavailable, genus-level assignments were used.

For each leaf on the tree, one representative genome was randomly selected. Gene annotations were derived from PADLOC^18,19^ and DefenseFinder^67^ for defence systems, the PFAM-A database^68^ scanned with pfam_scan.py and the geNomad database (v1.9)^62^ for domain annotations. Genomic contexts for each clade were visualized in R using ggplot2^69^ and gggenes. Detailed schemes with genomic neighbourhoods for each phylogenetic cluster are available in Zenodo.

To investigate the presence of RecBCD in Druantia III-containing genomes, we utilised HMM profiles from Mutte *et al* (20025)^70^ as queries in hmmsearch from HMMer^71^ package v 3.4 against all proteins in representative genomes.

### Protein conservation and positive selection

Conservation scores across full-length protein alignments were calculated using PyCanal^72^. To identify residues under positive selection in DruH, codon alignments were generated from protein alignments using PAL2NAL^73^, and corresponding phylogenetic trees were reconstructed with IQ-TREE^59^. Codon alignments and trees were used as input for HyPhy v2.5.93^51^. Sites under positive selection were inferred using both FUBAR^50^ and FEL^49^, and only residues supported by both methods were retained. For reference DruE/DruH pair, positively selected residues were mapped onto the AF3 model of the DruH WP_133301053.1 protein. For DruE WP_000213430.1, residues were mapped onto the AF3 model of DruH WP_001513100.1.

### Foldseek and FoldMason analysis of DruH similarity to phage baseplate J-like proteins

The full-length DruH structure was used as a query in Foldseek^74^ against the AFDB50 database to identify remote structural matches. Hits were sorted by probability and E-value, and those annotated as ‘baseplate protein J-like domain-containing proteins’ and an e-value < 1e-15 were retained. A representative subset of these top-scoring J-like hits was then analysed together with DruH using FoldMason^74^ to generate a structural multiple alignment. The FoldMason output was used to evaluate overall fold similarity across the selected proteins on the basis of structural superposition, guide tree, and alignment quality scores.

### Plasmid and strain construction

The plasmids constructed in this work are listed in **Table S5**, and the primers used can be found in **Table S6**. Plasmid pACYCDuet-1 was modified to contain the pBAD promoter from plasmid 8A (MacroLab) by Gibson assembly. YFP and Druantia III (from *E. coli* ECOR19) were cloned into the modified pACYCDuet-1 by Gibson assembly. YFP and Zorya II (ECOR19) were cloned into plasmid 8A by Gibson assembly. The phage genes were cloned into plasmids pCOLADuet-1 or pCDFDuet-1 by Gibson assembly. The DruE coding sequence was amplified by PCR and cloned into the pET21a vector to generate a C-terminal Strep-tagged fusion protein conferring ampicillin resistance. The DruH and ZorE genes were cloned independently into pET28 vectors to produce constructs carrying C-terminal Flag tags with kanamycin resistance. The plasmids were recovered in DH5α cells, extracted using Monarch® Spin Plasmid Miniprep Kit (New England Biolabs) and confirmed by sequencing at Plasmidsaurus (USA). Mutations of the genes were engineered by around-the-horn PCR and confirmed by Sanger sequencing (Macrogen). Plasmids were transformed individually or in combinations into competent BL21-AI cells prepared using the Mix&Go! *E. coli* Transformation Kit (Zymo).

### Efficiency of platting

Overnight cultures of the bacteria were diluted 1:50 in LB containing antibiotics, induced with 0.2% arabinose, and incubated for 4 hours before being used in double agar overlay assays. For this, bacterial cultures were mixed with 0.6% top agar and overlaid on LBA plates. Ten-fold serial dilutions of the phage stocks were spotted onto the bacterial lawn and the plates incubated overnight at 37 °C. The phage plaques were counted and used to calculate the EOP relative to the control. Statistical significance was determined using the multiple comparison function from Two-way ANOVA with a p-value of <0.01.

### Time post infection assay

Overnight bacterial cultures were diluted to an optical density at 600 nm of 0.1 in LB containing antibiotics and 0.2% arabinose. The cultures were infected with phage at an MOI of 10 or 0.1. At 0, 30, 60, 90 and 120 minutes post infection, a sample was taken and centrifuged at 12,000 × *g* for 2 minutes. The supernatant was serially diluted, and the phages were quantified by plaque assay on a bacterial lawn of cells with YFP. The pellets were washed twice, serially diluted and spotted on LBA plates. CFUs and PFUs were counted after overnight incubation at 37 °C.

### Liquid assay

Overnight bacterial cultures were diluted to an optical density at 600 nm of 0.1 in LB containing antibiotics and 0.2% arabinose. The bacterial suspension was distributed into wells of a 96-well plate to which phage dilutions or LB were added. The plates were incubated in a CLARIOstar Plus plate reader at 37 °C with shaking at 200 rpm, with optical density at 600 nm measured every 5 min for 12 h.

### Membrane permeability and depolarisation

Overnight cultures of control (YFP) and defence systems expressing cells were diluted to an OD at 600 nm of 0.1 in LB supplemented with antibiotics and 0.2% arabinose. The bacterial cultures were incubated at 37 °C, 200 rpm, until and OD600 of 0.4. The OD of the cultures were adjusted to 0.05 in LB supplemented with antibiotics, 0.2% arabinose, and 1% DMSO. For membrane permeability assays, PI was added to a final concentration of 5 µg/ml, while for membrane depolarisation, DISC3(5) was added to a final concentration of 10 nM. The cultures were incubated at 37 °C, 200 rpm, for 1 hour, and phages were then added at an MOI of 10 or 0.1. OD_600_, DISC(3)5 (excitation: 600 nm; emission: 670 nm), and PI (excitation: 544 nm; emission: 612 nm) were measured every 10 min.

### Generation of phage escape mutants

Phage plaques obtained in double layer agar plates with Druantia-Zorya cells were picked and dissolved in 30 µl of LB. The recovered phages were produced in liquid culture using the Druantia-Zorya-expressing strain as the host. The produced phages were serially diluted and spotted onto bacterial laws of Druantia-Zorya-expressing and control cells, to identify those phages able to escape Druantia-Zorya defence.

### Phage DNA extraction and sequencing

Phage DNA was extracted using the phenol-chloroform method. Briefly, DNase I and RNase were added to the phage stock at 1 µg/ml each, and the solution was incubated for 30 min. EDTA, proteinase K, and SDS were added to the solution at 20 mM, 50 µg/ml, and 0.5%, respectively, and incubated for 1 hour at 56 °C. An equal volume of phenol was added to the mixture, vortexed, and centrifuged at 3,000 × g for 10 min, at room temperature. The aqueous phase was recovered, mixed with an equal volume of 1:1 phenol:chloroform, and centrifuged. The procedure was repeated with chloroform. After centrifugation, the aqueous phase was recovered and mixed with 0.1 volumes of 3 M sodium acetate, pH 5.0 and 2.5 volumes of ice-cold absolute ethanol. The mixture was incubated at −20 °C for 1 hour and centrifuged at 14,000 × g for 15 min. The DNA pellet was resuspended in ice-cold 70% ethanol and centrifuged for 5 min. The pellet was let dry at room temperature and resuspended in nuclease-free water. The quality and quantity of extracted phage DNA were estimated using a Nanodrop and a Qubit fluorometer, respectively. Phage samples were sequenced at Novogene (China). Sample libraries were prepared using Novogene NGS DNA Library Prep Set, and sequenced on an Illumina platform, producing 1-2 Gb reads.

### Sequencing and analysis of phage escape mutants

For wild-type phage, seqtk version 1.3 (https://github.com/lh3/seqtk) was used to sample reads at 100x depth of the phage genomes. The subset reads were assembled using SPAdes version v3.15.351^75^ using default settings. For phage escape mutants, all reads were mapped to the assembled wild-type phage genome using bwa version 0.7.1752^76^ and samtools version 1.1153^77^. SNP calling was performed using gatk version 4.2.6.154^78^.

### ENDO-pore

ENDO-Pore experiments for DruE and supercoiled pUC19 were performed using the protocol previously described^40^, using 250 mM of protein and 1 µg of plasmid DNA, followed by incubation at 37 C for 30 min. The resulting Oxford Nanopore (ONT) reads were basecalled using Dorado (https://github.com/nanoporetech/dorado). Reads were quality-filtered using fastp^79^. After that we obtained 265,611 reads for DruE with median length of 4.2 kb. Consensus sequences from concatemeric reads were generated using C3POa^80^, with Cm used as the splint sequence. This resulted in 118,480 consensus sequences. The selected consensus sequences were aligned to pUC19 and Cm reference sequences using minimap2^81^ with the -ax map-ont preset. The resulting SAM files were processed with a custom Python script, which parsed CIGAR strings to reconstruct the arrangement of Cm and pUC19 segments within each consensus read. Reads of interest were defined as those containing Cm sequences at both ends and two internal alignments to pUC19: one spanning from the cleavage site to the end of the molecule, and the other from the start to the cleavage site. Cleavage positions were extracted from these alignments and visualised using a custom R script. To identify sequence contexts associated with cleavage, we extracted ±40 bp regions surrounding the most frequent cut sites and searched for palindromic motifs using MEME^82^ in oops mode.

### 3-aPBA assays

For efficiency of plating (EOP) assays, overnight bacterial cultures were diluted 1:50 in LB containing the appropriate antibiotics, 0.2% arabinose, and 3-aPBA at the indicated concentration, and incubated for 4 h at 37 °C before use in double agar overlay assays. For plating, bacterial cultures were mixed with 0.6% top agar containing the same concentration of 3-aPBA and overlaid onto LBA plates supplemented with the appropriate antibiotics, 0.2% arabinose, and 3-aPBA. Ten-fold serial dilutions of phage stocks were spotted onto the bacterial lawn and plates were incubated overnight at 37 °C. Plaques were counted and EOP was calculated relative to the appropriate control. For liquid infection assays, overnight cultures were diluted to OD_600_ of 0.1 in LB containing the appropriate antibiotics, 0.2% arabinose, and 3-aPBA at the indicated concentration. Cultures were distributed into 96-well plates, infected with phage at the indicated MOI, and incubated in a CLARIOstar Plus plate reader at 37 °C with 200 rpm shaking. OD_600_ was measured every 5 min for 12 h.

### Protein expression and purification

For protein expression, *E. coli* BL21 (DE3) cells were transformed with the appropriate expression plasmids and cultured in LB medium supplemented with the corresponding antibiotics at 37 °C until the OD_600_ reached 0.6-0.8. Protein expression was induced by the addition of 0.5 mM IPTG, followed by incubation at 18 °C for 16-18 h. Cells were harvested by centrifugation at 4,000 × g for 15 min at 4°C and stored at −80 °C until further purification.

For purification of Strep-tagged DruE wild-type protein and mutants, cell pellets were resuspended in lysis buffer containing 25 mM Tris-HCl (pH 8.0), 300 mM NaCl, 5% (v/v) glycerol, and 1 mM DTT, supplemented with SIGMAFAST Protease Inhibitor Cocktail Tablets, EDTA-Free, and lysed by sonication on ice. Cell lysates were clarified by centrifugation at 22,000 rpm for 1 h at 4 °C. The supernatant was incubated with Strep-Tactin TACS Agarose (IBA Lifesciences; 6-6350-010) pre-equilibrated in lysis buffer, with gentle rotation at 4 °C for 1 h. After extensive washing with lysis buffer, bound proteins were eluted using lysis buffer supplemented with 10 mM d-desthiobiotin (Sigma-Aldrich; D1411-1G). Eluted DruE-Strep proteins were further purified by size-exclusion chromatography (SEC) using a Superdex 200 Increase 10/300 GL column (GE Healthcare) equilibrated in SEC buffer containing 25 mM HEPES (pH 7.5), 150 mM NaCl, 0.2% (v/v) glycerol, and 1 mM DTT. Both wild-type and mutant DruE-Strep proteins eluted as homogeneous dimeric species. Peak fractions were pooled, concentrated to approximately 8 mg/ml, flash-frozen in liquid nitrogen, and stored at −80 °C.

For purification of Flag-tagged DruH or ZorE wild-type proteins and mutants, cell pellets were resuspended in the same lysis buffer described above and lysed by sonication. Cell lysates were clarified by centrifugation at 40,000 rpm for 1 h at 4 °C. The supernatant was incubated with anti-FLAG M2 affinity Gel (Sigma-Aldrich, A2220) for 3 h at 4 °C with gentle mixing. After extensive washing with lysis buffer, bound proteins were eluted using lysis buffer supplemented with 0.2 mg/ml 3×FLAG peptide (Sigma-Aldrich, F4799). Eluted proteins were concentrated and further purified by size-exclusion chromatography using a Superdex 200 Increase 10/300 GL column (GE Healthcare) equilibrated in SEC buffer containing 25 mM HEPES (pH 7.5), 150 mM NaCl, 0.2% (v/v) glycerol, and 1 mM DTT. DruH-Flag proteins typically eluted as a mixture of dimeric and monomeric species, whereas ZorE-FLAG eluted as a homogeneous monomer. Protein purity was assessed by SDS-PAGE.

### Cryo-EM sample preparation and data collection

Purified proteins were concentrated to 8 mg/ml. DNA substrates mixture with strand ratio 1:1 were annealed by heating to 95 °C, then gradually cooled to 25 °C in 5 °C steps, holding for 2 min at each step (15 bp duplex with a 15 nt 3′-OH overhang or a 15 bp duplex with a 28 nt 3′-OH overhang; sequences listed in **Table S6**). For DNA-bound samples containing the DruE dimer, DNA substrates were incubated with protein at a 1:1 molar ratio in the presence of 1 mM ATPγS and 2 mM MgCl_2_ for 30 min on ice. For the DruE-DruH-DNA sample, the components were incubated at a 1:1.2:1 molar ratio (DruE:DruH:DNA). For the DruH dimer ATP-bound structure, DruH dimers were incubated with a 28 nt 3’-OH DNA substrate in the presence of 1 mM ATP and 2 mM MgCl_2_ on ice for 30 min. Samples were then centrifuged at 12,000 rpm for 10 min prior to vitrification. To mitigate preferred particle orientation, fluorinated octyl maltoside (FOM) was freshly added to all samples immediately before freezing to a final concentration of 0.5 mM. A 4 µl aliquot of each sample was applied to glow-discharged holey carbon gold grids (Quantifoil Au 300 mesh R1.2/1.3). Grids were blotted for 2.5 s at 100% humidity and 6 °C and plunge-frozen into liquid ethane using a Vitrobot Mark IV (FEI). Grid screening was performed on a 300 kV Titan Krios G3 transmission electron microscope equipped with a K3 direct electron detector and operated using SerialEM software (Thermo Fisher Scientific).

For the DruE dimer apo, DruE dimer bound to a 28 nt 3’-OH DNA substrate, and DruH monomer apo samples, cryo-EM data were collected at the Memorial Sloan Kettering Cancer Center (MSKCC) using a Titan Krios G2 transmission electron microscope (FEI) operated at 300 kV and equipped with a K3 direct electron detector. Data acquisition was controlled using SerialEM software. Movies were recorded in super-resolution mode with a total electron dose of 53 e⁻ Å^-2^, a defocus range of −0.8 to −2.2 µm, and a calibrated physical pixel size of 1.064 Å.

For the DruE dimer bound to a 15 nt 3’-OH DNA substrate, DruE-DruH bound to a 28 nt 3’-OH DNA substrate, DruH dimer apo, and DruH dimer bound to ATP samples, cryo-EM data were collected at Memorial Sloan Kettering Cancer Center (MSKCC) using a Titan Krios G4 transmission electron microscope (FEI) operated at 300 kV and equipped with a Falcon 4i electron detector. Data collection was performed using EPU software. Movies were recorded in counting mode with 45 EER frames. The defocus range was set from −0.8 to −2.2 µm, with a calibrated pixel size of 0.725 Å and a total electron dose of 60 e⁻ Å^-2^.

### Cryo-EM data processing

For DruE dimer apo structure determination, a total of 3,885 movies were processed using cryoSPARC^83^ (v4.2.1). Patch-based motion correction and patch-based contrast transfer function (CTF) estimation were applied to correct beam-induced motion and estimate CTF parameters, respectively. Micrographs exhibiting ice contamination, high astigmatism, or poor CTF-fit resolution were excluded using the Manually Curate Exposures job with appropriate threshold criteria. The remaining high-quality micrographs were subjected to particle picking using the Blob Picker, followed by two-dimensional (2D) classification. Representative high-quality 2D classes were selected to generate templates for subsequent template-based particle picking. Particles were then extracted, yielding a total of 1,918,576 particles. Multiple rounds of 2D classification were performed to remove contaminants and poorly aligned particles, resulting in a final dataset of 207,272 selected particles. Ab initio reconstruction followed by heterogeneous refinement into three classes revealed two well-resolved classes corresponding to distinct symmetric and asymmetric dimer conformations. These two classes were selected for subsequent global and local CTF refinement to correct residual astigmatism and higher-order aberrations, followed by homogeneous refinement and non-uniform refinement. This processing yielded reconstructions at 4.04 Å resolution based on 54,773 particles for the symmetric dimer and 3.58 Å resolution based on 69,036 particles for the asymmetric dimer, respectively. The final density maps enabled reliable atomic model building for the asymmetric DruE dimer and main-chain tracing for the symmetric dimer, including well-resolved and consistent density at the dimeric interface in both conformational states.

For determination of the DruE dimer bound to a 15-nt 3’-OH DNA substrate, a total of 14,599 movies were processed using cryoSPARC^83^ (v4.2.1). Patch-based motion correction and CTF estimation were applied to correct beam-induced motion and estimate CTF parameters, respectively. Particles were initially picked using the Blob Picker and extracted, yielding a total of 2,965,052 particles. Multiple rounds of 2D classification were performed to remove contaminants and poorly aligned particles, resulting in a curated dataset of 742,569 particles. Following ab initio reconstruction and heterogeneous refinement into three classes, the best-resolved volume, containing 598,484 particles, was selected for further three-dimensional (3D) classification into six classes, revealing two distinct DNA-bound states: a symmetric and an asymmetric dimer. Two classes corresponding to the symmetric DNA-bound dimer were combined and subjected to homogeneous and non-uniform refinement, yielding a 3.23 Å resolution map based on 121,427 particles. The four classes corresponding to the asymmetric DNA-bound state were combined into a dataset of 477,057 particles and further classified into three 3D classes. The best-resolved class was selected for final homogeneous and non-uniform refinement, resulting in a 2.98 Å resolution map based on 154,317 particles. The resulting maps were of high quality and enabled reconstruction of the full-length DruE dimer bound to a 15-bp DNA duplex with a 15-nt 3’-OH overhang in both symmetric and asymmetric states, with well-defined ATPγS density observed in the helicase pocket of each DruE protomer.

To determine the structures of DruE complexes bound to a 28-nt 3’-OH DNA substrate, a total of 7,848 cryo-EM movies were processed using cryoSPARC^83^ (v4.2.1). Beam-induced motion was corrected by patch-based motion correction, and CTF parameters were estimated using patch-based CTF estimation. Initial particle picking was performed with the Blob Picker, followed by multiple rounds of 2D classification. Representative high-quality classes were selected to generate templates for subsequent template-based particle picking, after which particles were extracted, yielding 691,330 particles. Two distinct particle populations were identified during 2D classification, corresponding to DruE dimers bound to DNA (443,544 particles) and DruE trimers bound to DNA (73,848 particles). These datasets were independently subjected to ab initio reconstruction and heterogeneous refinement to further eliminate contaminants and poorly aligned particles. Within the dimeric population, symmetric and asymmetric DNA-bound conformations were resolved, comprising 125,395 and 220,818 particles, respectively. Particles corresponding to the symmetric dimer were further refined by homogeneous refinement and 3D classification into four classes to enhance map quality. Two well-resolved classes, containing a combined total of 65,695 particles, were selected for global and local CTF refinement to improve optical parameter estimation, followed by final homogeneous and non-uniform refinement. This workflow yielded a high-quality reconstruction of the symmetric DruE dimer bound to a 15-bp DNA duplex with a 28-nt 3’-OH overhang, enabling subsequent model building. The asymmetric dimer and trimer datasets were processed using analogous refinement strategies, including global and local CTF refinement and final homogeneous and non-uniform refinement. These procedures resulted in reconstructions at 3.18 Å resolution for the asymmetric dimer (220,818 particles) and 3.72 Å resolution for the trimeric assembly (64,246 particles), respectively. The resulting density maps supported reliable atomic modeling of the full-length DruE dimer and trimer, with well-resolved density corresponding to a 15-bp duplex DNA with either an 18-nt or 15-nt 3’-OH overhang. In all structures, clear density for ATPγS was observed within the helicase pocket of each DruE protomer. Notably, the trimeric assembly revealed two additional ATPγS molecules located at the interfaces between the third DruE protomer and the preformed DruE dimer, mediated by interactions involving the C-terminal domains, suggesting that this region may represent a potential DruE nuclease pocket.

For determination of the DruE-DruH-DNA complex, movie stacks were first subjected to patch motion correction and patch CTF estimation. Micrographs with poor CTF fits or significant drift were discarded. Particles were automatically picked using both blob picker and template picker, followed by several rounds of 2D classification to remove false positives and poorly aligned particles. Particles corresponding to dimeric DruE and monomeric DruE were then processed separately and subjected to ab initio reconstruction followed by heterogeneous refinement into two classes to remove junk particles. This classification step separated particles corresponding to monomeric DruE–DNA complexes, dimeric DruE particles, and DruH monomer assemblies present in the dataset. A total of 278,600 particles belonging to the monomeric DruE–DNA class were selected for further refinement. The selected particles were refined using homogeneous refinement, followed by global and local CTF refinement and non-uniform refinement to improve map quality. The final reconstruction yielded a cryo-EM density map of monomeric DruE bound to the DNA substrate at 2.97 Å resolution with 192,121 particles, with clear density observed for both the duplex DNA region and the single-stranded 3’ overhang.

For determination of the DruH apo structures, 4,539 movies (monomer dataset) and 6,165 movies (dimer dataset) were collected and processed using cryoSPARC^83^ (v4.2.1). For the DruH monomer apo dataset, particle picking was performed using both Blob Picker and Template Picker. A total of 1,427,621 particles (binned by 2) were extracted and subjected to multiple rounds of 2D classification to remove junk particles and poorly aligned classes. After curation, 1,084,353 particles were selected for 3D reconstruction, including ab-initio reconstruction and heterogeneous refinement into three classes. Two identical high-quality classes comprising 909,860 particles were combined, re-extracted at bin1, and further refined through ab-initio reconstruction and heterogeneous refinement into two classes. The best-resolved class, containing 684,009 particles, was selected for global and local CTF refinement, followed by homogeneous and non-uniform refinement, yielding a final reconstruction at 3.49 Å resolution.

For the DruH dimer apo dataset, particle picking using Blob and Template Pickers yielded 1,032,681 particles, which were subjected to multiple rounds of 2D classification. After curation, 501,375 high-quality particles were retained for ab-initio reconstruction and heterogeneous refinement into four classes. One well-resolved class containing 80,084 particles was selected for global and local CTF refinement, followed by homogeneous and non-uniform refinement, resulting in a DruH dimer structure (state 1) at 3.66 Å resolution. Another heterogeneous refinement class, consisting of 100,558 particles, represented a mixture of conformational states and was further subjected to additional 3D classification into three classes. Two of these classes were independently refined through global and local CTF refinement, homogeneous refinement, and non-uniform refinement, yielding DruH dimer state 2 at 4.41 Å resolution (27,338 particles) and DruH dimer state 3 at 4.47 Å resolution (14,396 particles), respectively. Subsequent model building enabled reliable atomic modeling of the DruH monomer and the dimer state 1. For dimer states 2 and 3, main-chain tracing was achieved. In all reconstructions, the density maps were consistent with well-defined α-helical and β-sheet features in the monomer and clearly resolved dimerization interfaces across the three apo dimer states.

For determination of the DruH dimer bound to ATP and Mg^2+^, a total of 11,491 movies were collected and processed using cryoSPARC^83^ (v4.2.1). After standard image preprocessing, including patch-based motion correction and CTF estimation, micrographs were selected for subsequent particle picking. Initial particle picking was performed using the Blob Picker, followed by 2D classification to generate representative templates for template-based particle picking. Using the Template Picker, 929,336 particles were selected and extracted from the micrographs. These particles were subjected to two additional rounds of 2D classification to remove contaminants and poorly aligned particles, resulting in a curated dataset of 614,466 particles. The selected particles were then subjected to ab-initio reconstruction and heterogeneous refinement into three classes. One well-resolved class was selected and further refined through 3D classification into four classes to improve separation of intact volumes. Two highly similar and well-defined classes, comprising a combined total of 83,407 particles, were merged and subjected to homogeneous refinement using the class 0 volume as the initial model. Global and local CTF refinement were subsequently applied, followed by non-uniform refinement, yielding a final reconstruction at 3.54 Å resolution. The resulting density map enabled accurate atomic model building of the full-length DruH dimer, with well-resolved density for ATP and Mg^2+^ observed in the active site of each DruH protomer. The dimeric interface was also clearly defined, with density features in strong agreement with the refined atomic model.

### Model building and refinement

Initial atomic models were generated using structure predictions from AlphaFold3^84^ and rigid-body docked into the corresponding cryo-EM density maps. Manual model adjustment was performed in Coot^85^ to correct side-chain conformations, rebuild flexible loops, and position ligands and Mg^2+^ ions according to the density. Real-space refinement was carried out in PHENIX^86^ with appropriate stereochemical and geometry restraints applied throughout the refinement process. Model quality was evaluated using standard validation metrics, including Ramachandran statistics, rotamer analysis, and map-to-model correlation coefficients.

### Purification of tagless DruH and ZorE proteins

For preparation of tagless proteins, DruH and ZorE were cloned into a pET-based expression vector containing an N-terminal Flag tag followed by a TEV protease cleavage site and transformed into *E. coli* BL21 (DE3) cells for recombinant expression. Cells were grown in LB medium supplemented with the appropriate antibiotic at 37 °C to an OD_600_ of ∼0.6-0.8, followed by induction with 0.5 mM IPTG and overnight expression at 16 °C. Protein purification was performed following the same protocol as for DruH-Flag. Differently, after elution from Flag affinity resin, the protein was incubated with TEV protease at 4 °C overnight to remove the N-terminal Flag tag. Cleavage efficiency was assessed by SDS-PAGE. Upon complete tag removal, the sample was passed over Ni-NTA resin to remove His-tagged TEV protease. The flow-through, containing tagless DruH or ZorE, was collected and concentrated. The protein was further purified by SEC on a Superdex 200 Increase column equilibrated in gel filtration buffer. Peak fractions corresponding to tagless DruH (dimeric and monomeric species) or ZorE were pooled, concentrated, flash-frozen in liquid nitrogen, and stored at −80 °C.

### Mass photometry

Mass photometry measurements were carried out using a Refeyn TwoMP instrument. A pre-assembled 6-well gasket (Refeyn) was placed onto a clean sample carrier slide (Refeyn), with each well used for a separate measurement. To define the focal plane, 15 μl of freshly prepared SEC buffer (25 mM HEPES, pH 7.5, 150 mM NaCl, 0.2% (v/v) glycerol) or DNA-ATPγS-Mg^2+^ supplemented buffer (SEC buffer supplemented with 2 mM MgCl_2,_ 1 mM ATPγS, and 1 μM DNA) was added to each well as a reference. The focal position was established and maintained throughout acquisition using the instrument’s built-in autofocus system based on total internal reflection. Purified N-Flag-DruE dimer, tagless DruH dimer, tagless DruH monomer, and tagless ZorE monomer were analysed either individually or in combination, in the presence or absence of DNA (1 μM freshly annealed 15 bp duplex with a 28-nt 3’ overhang), 1 mM ATPγS, and 2 mM MgCl_2_. Samples were prepared in the same buffer used for purification and incubated on ice for 30 min prior to measurement. Proteins were first diluted to 200 nM, and 3 μl of sample was introduced into the buffer drop within the measurement well, resulting in a final concentration of 33.3 nM. Following autofocus stabilisation, movies were collected for 60 s. Data were acquired using Refeyn AcquireMP (v2024.1.1.0) and processed with Refeyn DiscoverMP (v2024.1.0.0). Calibration of contrast-to-mass conversion was performed using bovine serum albumin (BSA; Sigma) as a reference standard. Mass distributions were analysed in DiscoverMP by fitting Gaussian functions to determine the mean molecular mass of each species. Final plots were generated using GraphPad Prism 11.

### Protein purification for pull-down assays

N-terminally Flag-tagged DruE was cloned into a pET21a vector with an intervening GGS linker between the TEV site and DruE, using ampicillin selection, and expressed in *E. coli* BL21 (DE3) cells. Protein expression and affinity purification followed the same procedure as described for C-terminally Flag-tagged DruH. Eluted protein was further purified by SEC using a Superose 200 Increase 10/300 GL column equilibrated in buffer containing 25 mM HEPES (pH 7.5), 150 mM NaCl, 0.2% glycerol, and 2 mM DTT. Peak fractions were pooled and concentrated.

GamS with a C-terminal twin Strep II tag was cloned into a pET21a vector with ampicillin selection and expressed in *E. coli* BL21 (DE3) cells. Protein expression and purification followed the same procedure as described for C-terminally Strep-tagged DruE. The eluate was further purified by SEC using a Superdex 75 Increase 10/300 GL column equilibrated in buffer containing 25 mM HEPES (pH 7.5), 150 mM NaCl, 0.2% glycerol, and 2 mM DTT. Peak fractions were pooled and concentrated.

Ocr with a C-terminal His_6_ tag was cloned into a pYB100 vector with kanamycin selection and expressed in *E. coli*BL21 (DE3) cells. Protein expression was induced with 0.5 mM IPTG at 16 °C for ∼20 h. Cells were resuspended in lysis buffer (25 mM Tris-HCl pH 8.0, 500 mM NaCl, 5% glycerol, 2 mM β-mercaptoethanol, supplemented with PMSF) and lysed by sonication. Lysates were clarified by centrifugation (22,000 rpm, 1 h, 4 °C) and applied to Ni-NTA resin. After washing, protein was eluted with buffer containing 500 mM imidazole. The eluate was further purified by SEC using a Superdex 75 Increase column equilibrated in buffer containing 25 mM HEPES (pH 7.5), 150 mM NaCl, 5% glycerol, and 2 mM DTT. Peak fractions were pooled and concentrated.

### *In vitro* pull-down assays

Strep-tagged GamS or His-tagged Ocr was mixed with Flag-tagged DruE, DruH (monomeric or dimeric fractions), or ZorE at approximately equimolar ratios in 500 µl binding buffer (25 mM HEPES pH 7.5, 150 mM NaCl, 0.2% glycerol, 1 mM DTT, 0.2% Triton X-100). Samples were incubated at 4 °C for 30 min with gentle rotation. An aliquot (40 µl) was removed as input prior to immunoprecipitation. The mixtures were then incubated with 50 µl anti-FLAG M2 affinity resin for 2.5 h at 4 °C with gentle rotation. The resin was washed five times with binding buffer. Bound proteins were eluted with binding buffer containing 0.2 mg/mL 3×FLAG peptide by incubation on ice for 30 min with gentle mixing every 5 min. Input and elution fractions were analysed by SDS-PAGE and transferred onto PVDF membranes using a semi-dry transfer system (Bio-Rad). Membranes were blocked with 5% (w/v) nonfat dry milk for 1 h at room temperature and incubated overnight at 4 °C with HRP-conjugated anti-Flag (Cell Signaling, 86861S), anti-His (Thermo Fisher, MA1-21315-HRP), or anti-Strep II (Abcam, ab324294) antibodies as indicated (1:4,000 dilution). After washing with TBS containing 0.05% Tween-20, signals were detected using enhanced chemiluminescence (Thermo Scientific) and visualised on a ChemiDoc MP imaging system (Bio-Rad).

### Co-immunoprecipitation of Bas37_0273 and DruH

For co-IP assays, Bas37_0273 and Bas37_0273Q284R were individually cloned into the PYB100-Kan backbone with a C-terminal His tag. The resulting His-tagged constructs were each co-transformed into BL21(DE3) cells together with N-Flag-DruH cloned in the pET21a vector. As negative controls, Bas37_0273-His or Bas37_0273Q284R-His alone were transformed into BL21 (DE3) cells. For each sample, cells were grown in 1 L of LB medium supplemented with the appropriate antibiotics and protein expression was induced with 0.5 mM IPTG. Cell pellets were harvested and resuspended in 20 ml lysis buffer (150 mM NaCl, 25 mM HEPES pH 7.5, 2 mM DTT, and 2% (v/v) glycerol), supplemented with 0.2% Triton X-100 to reduce nonspecific binding. Cells were lysed by sonication, and lysates were clarified by centrifugation at 22,000 rpm for 50 min. The supernatants were collected for subsequent co-IP analysis. To prepare input samples, 45 μl of each clarified supernatant were mixed with 15 μl of 4× SDS loading buffer. The remaining supernatant was incubated with 50 μl anti-Flag resin at 4 °C for 2.5 h with gentle rotation. After binding, the resin was washed five times with lysis buffer containing 0.2% Triton X-100. Bound proteins were eluted using 200 mM 3×Flag peptide in elution buffer consisting of lysis buffer supplemented with 3×Flag peptide. Elution was carried out on ice for 30 min, with gentle mixing every 5 min. Input and elution fractions were analysed by SDS-PAGE and transferred onto PVDF membranes using a semi-dry transfer system (Bio-Rad). Membranes were blocked with 5% (w/v) nonfat dry milk for 1 h at room temperature and then incubated overnight at 4 °C with HRP-conjugated anti-Flag antibody (Cell Signaling, 86861S) or HRP-conjugated anti-His antibody (Thermo Fisher, MA1-21315-HRP) at a 1:4,000 dilution, as indicated. After washing with TBS containing 0.05% Tween-20, signals were detected using enhanced chemiluminescence (Thermo Scientific) and visualised on a ChemiDoc MP imaging system (Bio-Rad).

### Structural alignment, analysis, and visualization

Structural superposition and root-mean-square deviation (RMSD) calculations were performed using UCSF ChimeraX^87^. Structural representations and figures were generated using UCSF ChimeraX and UCSF Chimera, and composite panels were assembled in Adobe Photoshop and Adobe Illustrator.

### Quantification and statistical analysis

Unless stated otherwise, experiments were performed in biological triplicates and are represented as the mean and standard deviation. Statistical significance was calculated by ratio paired t-test or by two-way ANOVA with sidak’s multiple comparison test, with a significance level of 0.05.

## Data availability

All unique bacterial strains, phages, and plasmids generated in this study are available from the lead contact without restriction. Raw data have been deposited at the Protein Data Base (PDB 11PQ, 11PT, 11PZ, 11QB, 11TO, 11WJ, 11WU, 11WW, 11WY, 11XD, 11XF, 11XO, 11XQ) and the Electron Microscopy Data Bank (EMDB EMD-75932, EMD-75936, EMD-75937, EMD-75943, EMD-76041, EMD-76129, EMD-76141, EMD-76145, EMD-76147, EMD-76152, EMD-76153, EMD-76162, EMD-76164) and are publicly available as of the date of publication. All original code used for data analysis has been deposited in Zenodo at https://zenodo.org/records/19900874.

