## Supplementary material for "Antiphage defence systems Druantia III and Zorya II synergise via shared DNA intermediates in a phage-specific manner": Figure S

### Contents

**Figure S1.** Phylogenetic, ecological, and genomic-context analysis of Druantia III and associated Zorya II components.

**Figure S2.** DNA mimic proteins Ocr and GamS do not form detectable stable complexes with DruE, DruH, or ZorE under the conditions tested.

**Figure S3.** Structural comparison of symmetric and asymmetric DruE dimers.

**Figure S4.** Conserved SF2 helicase motifs, zinc-binding clusters, and conformational changes in DNA-bound DruE.

**Figure S5.** Additional DruE assemblies and DNA-processing features.

**Figure S6.** Additional evolutionary, biophysical, and structural analysis of DruH.

**Figure S7.** Additional analyses of ZorE assemblies.

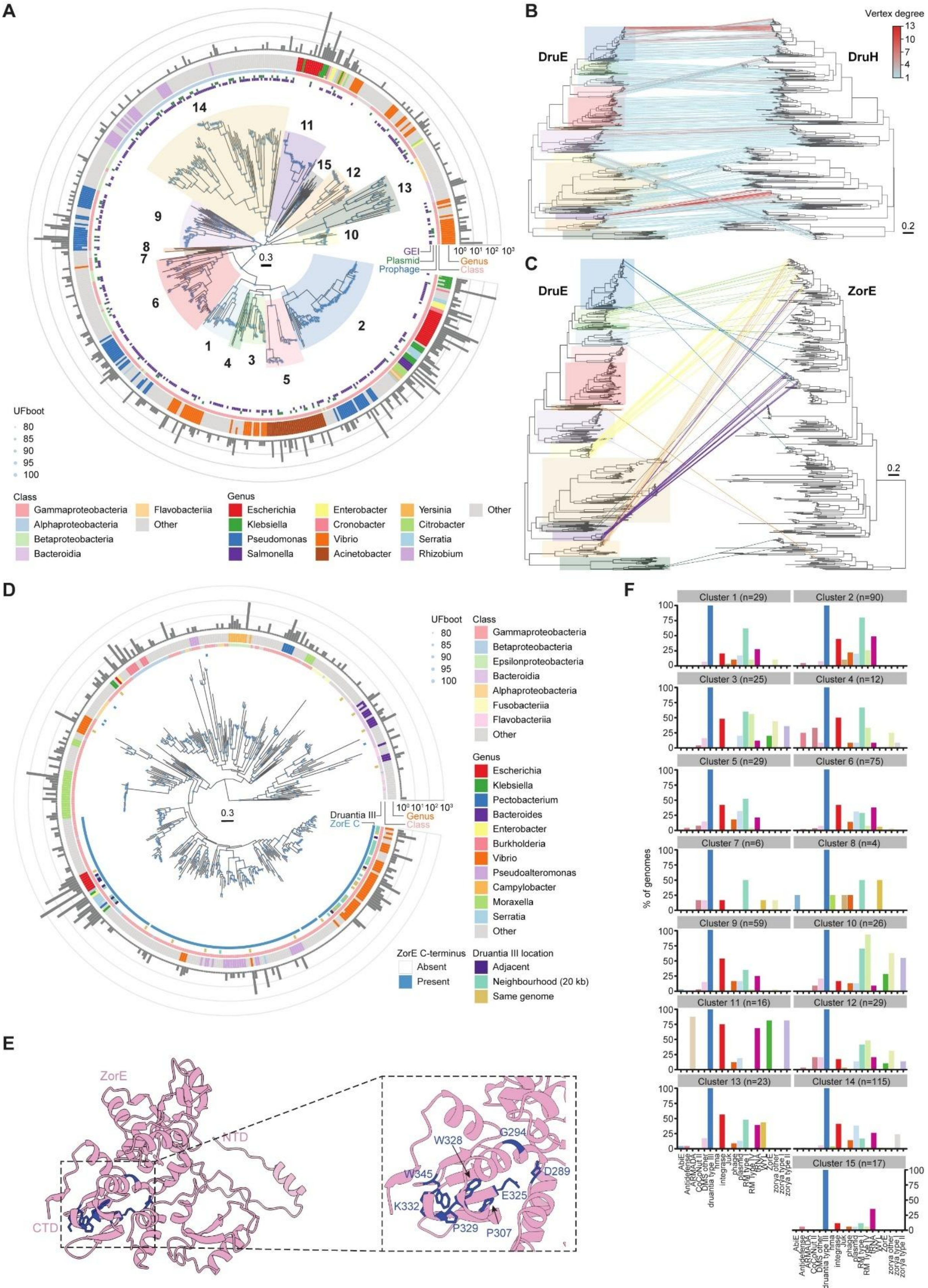


**Figure S1.** Phylogenetic, ecological, and genomic-context analysis of Druantia III and associated Zorya II components. **(A)** Maximum-likelihood phylogeny of dereplicated DruH proteins from complete Druantia III systems. The 15 DruH clusters are indicated. Node support is shown by ultrafast bootstrap (UFboot) values, and the outer tracks indicate taxonomic assignment and genomic context, including association with genomic islands (GEIs), plasmids and prophages. Bar plots on the outside indicate the number of representatives in each lineage. Phylogenetic clusters are shown as background segments, with colours maintained consistently across all panels. **(B)** Tanglegram linking the phylogenies of DruE and DruH proteins. Lines connect cognate DruE-DruH pairs, and line colour indicates vertex degree, highlighting the extent of partner connectivity across the dataset. Shaded boxes indicate the major DruE clusters. **(C)** Full tanglegram linking the phylogenies of DruE and ZorE. Lines connect cognate DruE-ZorE pairs, and are coloured according to DruE phylogenetic clusters, highlighting the extent of partner connectivity across the dataset. Shaded boxes indicate the major DruE clusters. **(D)** Maximum-likelihood phylogeny of dereplicated ZorE proteins from Zorya type II systems. Node support is shown by ultrafast bootstrap (UFboot) values, and the outer tracks indicate taxonomic assignment, association with Druantia III (adjacent, within a 20-kb neighbourhood, or elsewhere in the same genome), and the presence or absence of the ZorE C-terminal extension. Bar plots on the outside indicate the number of representatives corresponding to each leaf. **(E)** AlphaFold 3 (AF3) predicted structure of ZorE highlighting conserved residues within the nuclease-like C-terminal region. Left, overall structure of ZorE with the N-terminal domain (NTD) and C-terminal domain (CTD) indicated. Right, close-up view of the boxed region, showing conserved C-terminal residues as blue sticks. **(F)** Genomic neighbourhood composition surrounding Druantia III loci across the 15 DruE clusters. Bars indicate the percentage of genomes in each cluster containing the indicated gene categories within a 20-kb region centred on the Druantia III locus. Additional information with 20 kb neighbourhood visualisations is provided in Table S2 and in the associated dataset (see Key Resources Table).

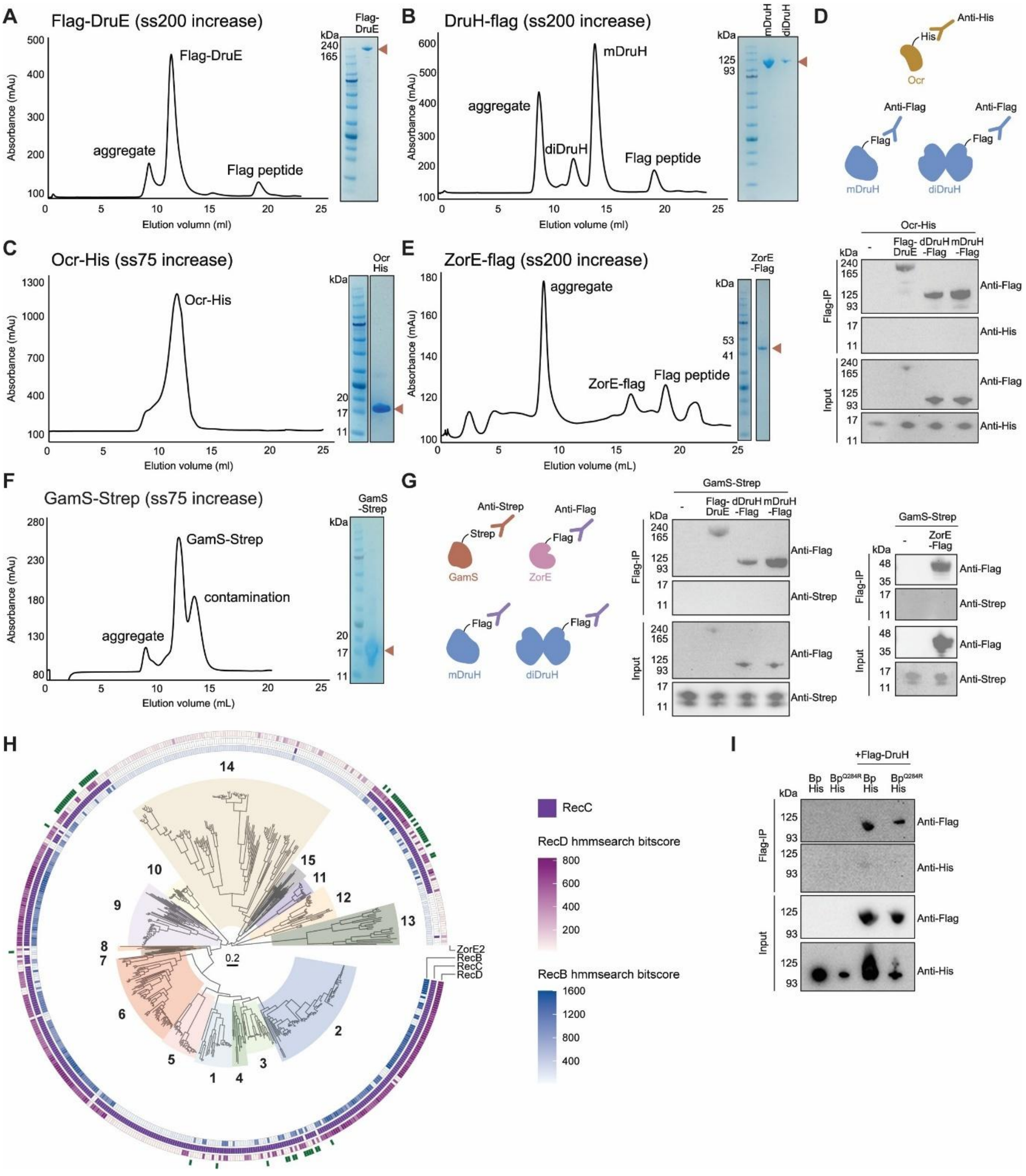


**Figure S2.** DNA mimic proteins Ocr and GamS do not form detectable stable complexes with DruE, DruH, or ZorE under the conditions tested. **(A)** Size-exclusion chromatography (SEC) profile of Flag-DruE on a Superdex 200 Increase column, with the corresponding Coomassie-stained SDS-PAGE gel of the peak fraction. Peaks corresponding to aggregate, Flag-DruE, and free Flag peptide are indicated. **(B)** SEC profile of DruH-Flag on a Superdex 200 Increase column, showing aggregate, dimeric DruH (diDruH), monomeric DruH (mDruH), and free Flag peptide. Right, corresponding Coomassie-stained SDS-PAGE gel of the monomeric and dimeric DruH peak fractions. **(C)** SEC profile of Ocr-His on a Superdex 75 Increase column, with the corresponding Coomassie-stained SDS-PAGE gel of the peak fraction on the right.

**(D)** Co-immunoprecipitation assays testing interaction of Ocr-His with Flag-DruE, dimeric DruH-Flag, or monomeric DruH-Flag. Schematics of the proteins and tags used are shown above. Anti-Flag immunoprecipitation was followed by immunoblotting with anti-Flag and anti-His antibodies. **(E)** SEC profile of ZorE-Flag on a Superdex 200 Increase column, showing aggregate, ZorE-Flag, and free Flag peptide, with the corresponding Coomassie-stained SDS-PAGE gel of the peak fraction shown on the right. **(F)** SEC profile of GamS-Strep on a Superdex 75 Increase column, with the corresponding Coomassie-stained SDS-PAGE gel of the peak fraction. **(G)** Co-immunoprecipitation assays testing interaction of GamS-Strep with ZorE-Flag, Flag-DruE, dimeric DruH-Flag, or monomeric DruH-Flag. Schematics of the proteins and tags used are shown on the left. Anti-Flag immunoprecipitation was followed by immunoblotting with anti-Flag and anti-Strep antibodies. (**H**) Distribution of RecBCD components and ZorE across representative Druantia III-containing genomes. Maximum-likelihood phylogeny of dereplicated DruE proteins from complete Druantia III systems, with one randomly selected representative genome shown per leaf. Branch sectors are coloured by DruE cluster. Concentric tracks indicate the presence of RecBCD components and ZorE in the corresponding genomes. **(I)** Co-immunoprecipitation assays testing interaction of Bas37_0273-His (Bp-His) and Bas37_0273- Q284R-His ($Bp^{Q284R}$-His) with DruH-Flag. Anti-Flag immunoprecipitation was followed by immunoblotting with anti-Flag and anti-His antibodies.

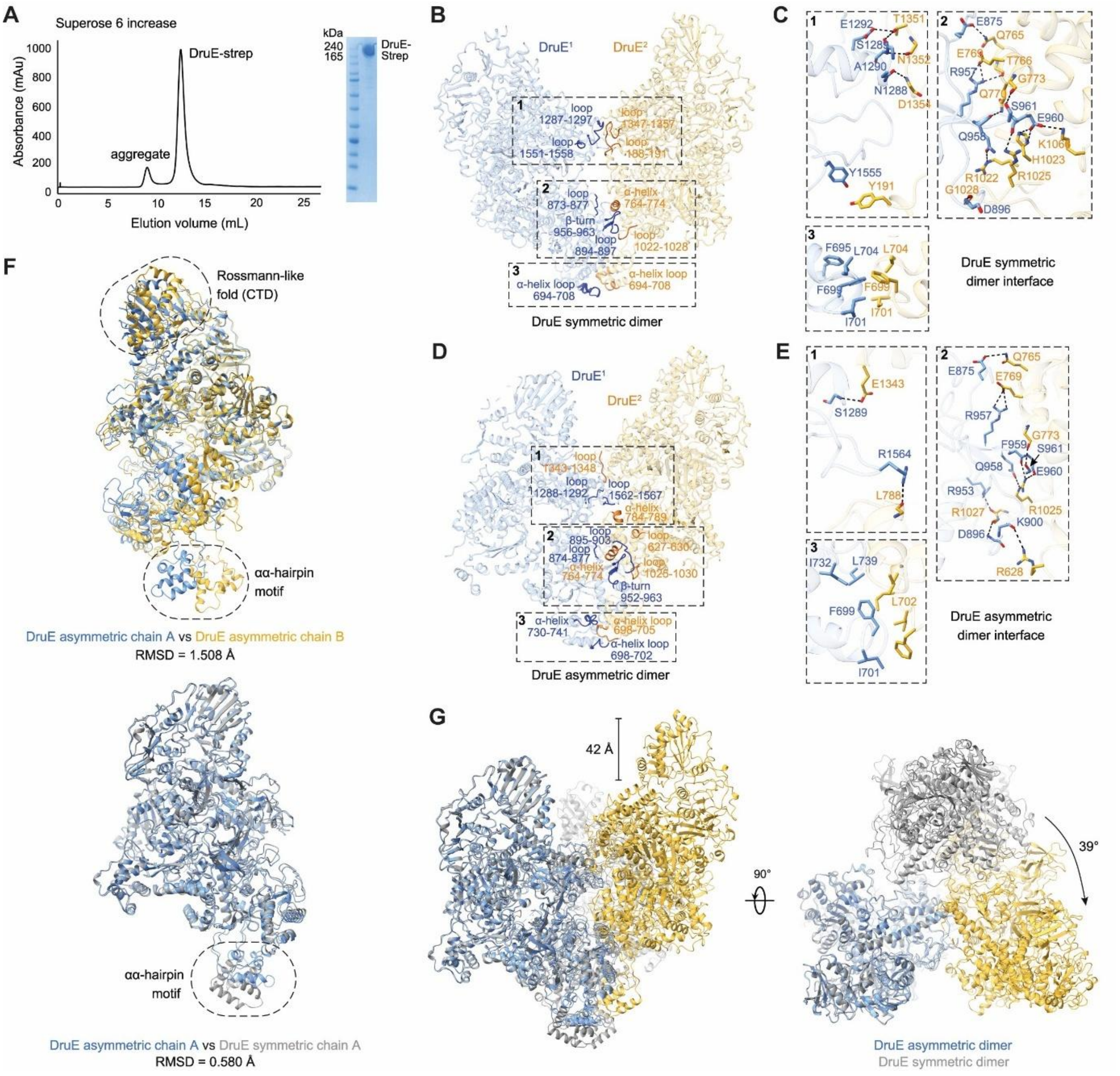


**Figure S3.** Structural comparison of symmetric and asymmetric DruE dimers. **(A)** Size-exclusion chromatography profile of Strep-tagged DruE on a Superose 6 Increase column, with the corresponding Coomassie-stained SDS-PAGE gel of the peak fraction. Peaks corresponding to aggregate and DruE-Strep are indicated. **(B)** Overall structure of the apo DruE symmetric dimer. The two protomers are coloured blue and gold. Dashed boxes indicate three regions contributing to the dimer interface. **(C)** Close-up views of the three interface regions highlighted in (B), showing residues that stabilise the symmetric dimer interface. **(D)** Overall structure of the apo DruE asymmetric dimer. The two protomers are coloured blue and gold. Dashed boxes indicate three regions contributing to the dimer interface. **(E)** Close-up views of the three interface regions highlighted in (D), showing residues that stabilise the asymmetric dimer interface. **(F)** Structural superpositions highlighting conformational differences within and between DruE dimers. Top, superposition of DruE asymmetric chain A and chain B (RMSD = 1.508 Å), highlighting displacement of the Rossmann-like fold in the C-terminal domain (CTD) and the αα-hairpin motif. Bottom, superposition of DruE asymmetric chain A and DruE symmetric chain A (RMSD = 0.580 Å), showing the close overall similarity between protomers from the two assemblies. **(G)** Structural comparison of the symmetric and asymmetric DruE dimers, showing conformational differences between the two assemblies, including a displacement of approximately 42 Å and a relative rotation of approximately 39° between corresponding protomers. The right panel shows the same comparison after a 90° rotation.

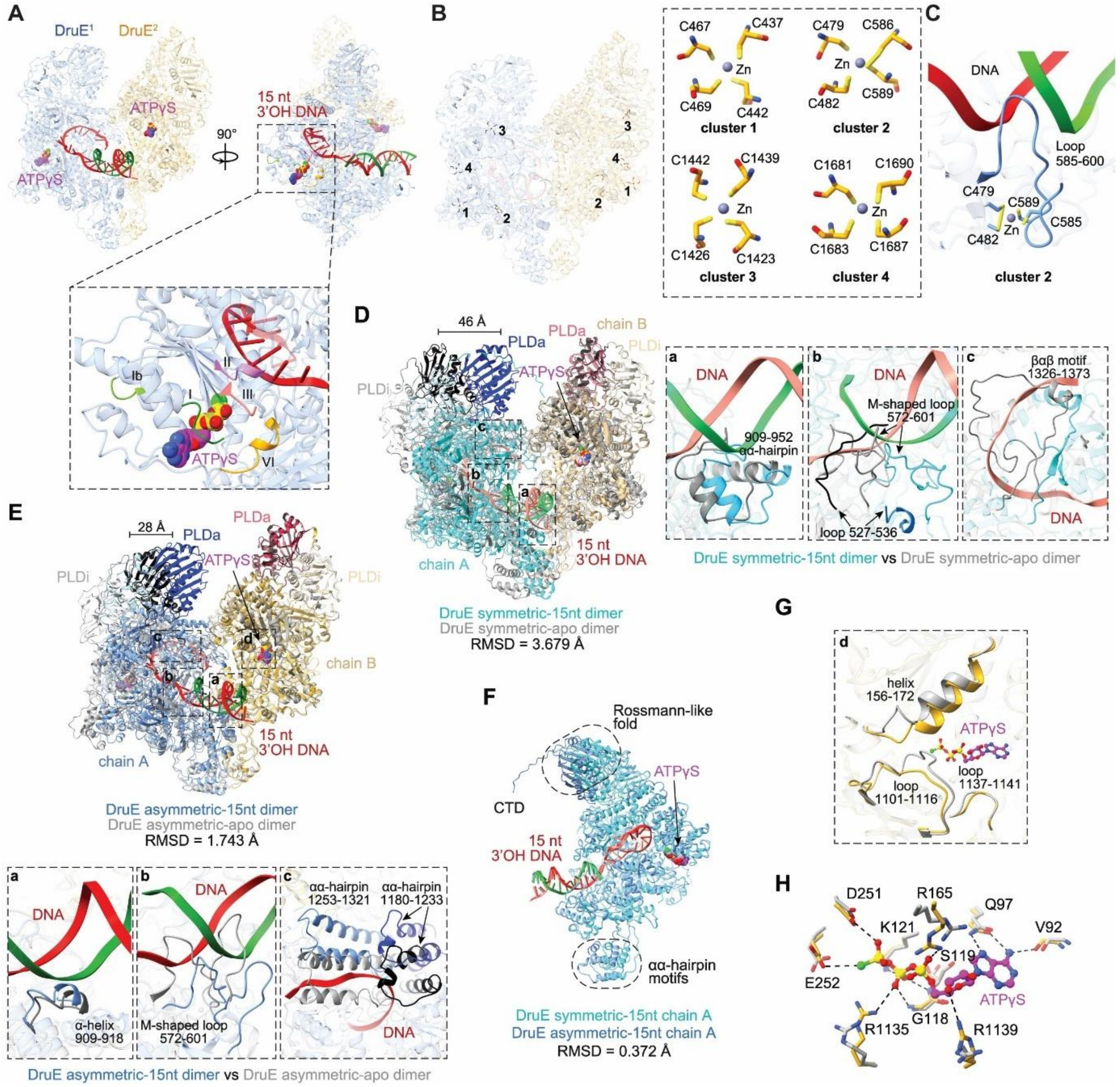


**Figure S4.** Conserved SF2 helicase motifs, zinc-binding clusters, and conformational changes in DNA-bound DruE. **(A)** Overall structure of the DruE asymmetric dimer bound to a 15-bp duplex DNA containing a 15-nt 3' overhang in the presence of ATPγS. The two protomers are coloured blue and gold. DNA is shown in red and green, and ATPγS molecules are shown in magenta. Insets highlight the conserved helicase motifs Ib, II, III and VI surrounding the ATPγS-binding pocket. **(B)** Positions of the four zinc-binding clusters in the DruE dimer. **(C)** Close-up view of zinc-binding cluster 2 located adjacent to the DNA-binding region, showing the coordinating cysteine residues and its position relative to the DNA substrate and loop 585–600. **(D)** Structural comparison of the symmetric DruE dimer bound to 15-nt 3'-overhang DNA with the symmetric apo dimer (grey) (RMSD = 3.679 Å). Insets highlight conformational changes in the αα-hairpin region (residues 909-952), the M-shaped loop (residues 572-601), the βαβ motif (residues 1326-1373), and the C-terminal region containing the PLD domains. **(E)** Structural comparison of the asymmetric DruE dimer bound to 15-nt 3'-overhang DNA with the asymmetric apo dimer (grey) (RMSD = 1.743 Å). Insets highlight conformational changes in the α-helix 909-918, the M-shaped loop (residues 572-601), the αα-hairpin motifs (residues 1180-1233 and 1253-1321), and the C-terminal region containing the PLD domains. **(F)** Superposition of chain A from the symmetric and asymmetric DNA-bound DruE dimers (RMSD = 0.372 Å), highlighting the relative movement of the Rossmann-like fold in the C-terminal domain (CTD) and the αα-hairpin motifs. **(G)** Close-up view of the nucleotide-binding region, showing conformational changes in helix 156-172 and surrounding loops near the ATPγS-binding pocket. **(H)** Detailed interactions between ATPγS and surrounding residues within the helicase pocket.

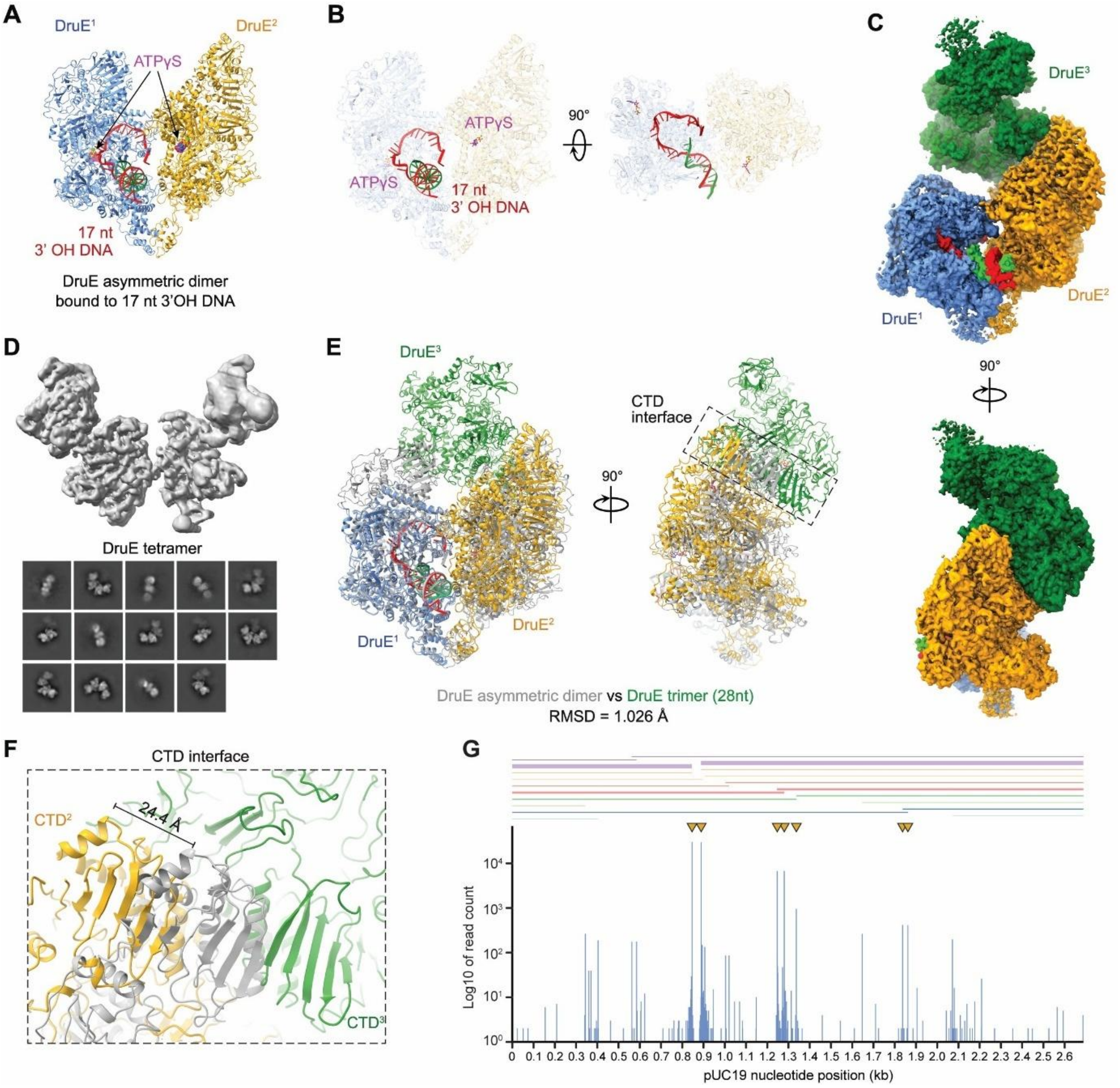


**Figure S5.** Additional DruE assemblies and DNA-processing features. **(A)** Cryo-EM structure of the asymmetric DruE dimer bound to a 15-bp duplex DNA containing a 17-nt 3′ overhang in the presence of ATPγS. The two protomers are coloured blue and gold. **(B)** Two views of the DNA trajectory in the asymmetric DruE dimer shown in (A), highlighting the path of the extended 3' single-stranded region through the dimer. **(C)** Cryo-EM density map of a trimeric DruE assembly observed in the 28-nt overhang dataset. The three protomers are coloured blue, gold, and green. **(D)** Low-resolution cryo-EM density map and representative 2D class averages of a tetrameric DruE assembly. **(E)** Structural comparison of the asymmetric DruE dimer and the DruE trimer from the 28-nt overhang dataset (RMSD = 1.026 Å). Superposition highlights the position of the third DruE protomer and the CTD-mediated interface. **(F)** Close-up view of the CTD interface formed between the second and third DruE protomers in the trimer, showing the approximately 24.4 Å repositioning of the CTD associated with trimer formation. **(G)** ENDO-pore cleavage profile of DruE on supercoiled pUC19 DNA. Blue bars indicate cleavage frequency across the plasmid, and yellow triangles mark recurrent cleavage hotspots. The aligned sequence regions corresponding to the 10 most common states, with line width indicating the relative frequency it is observed in the experiment.

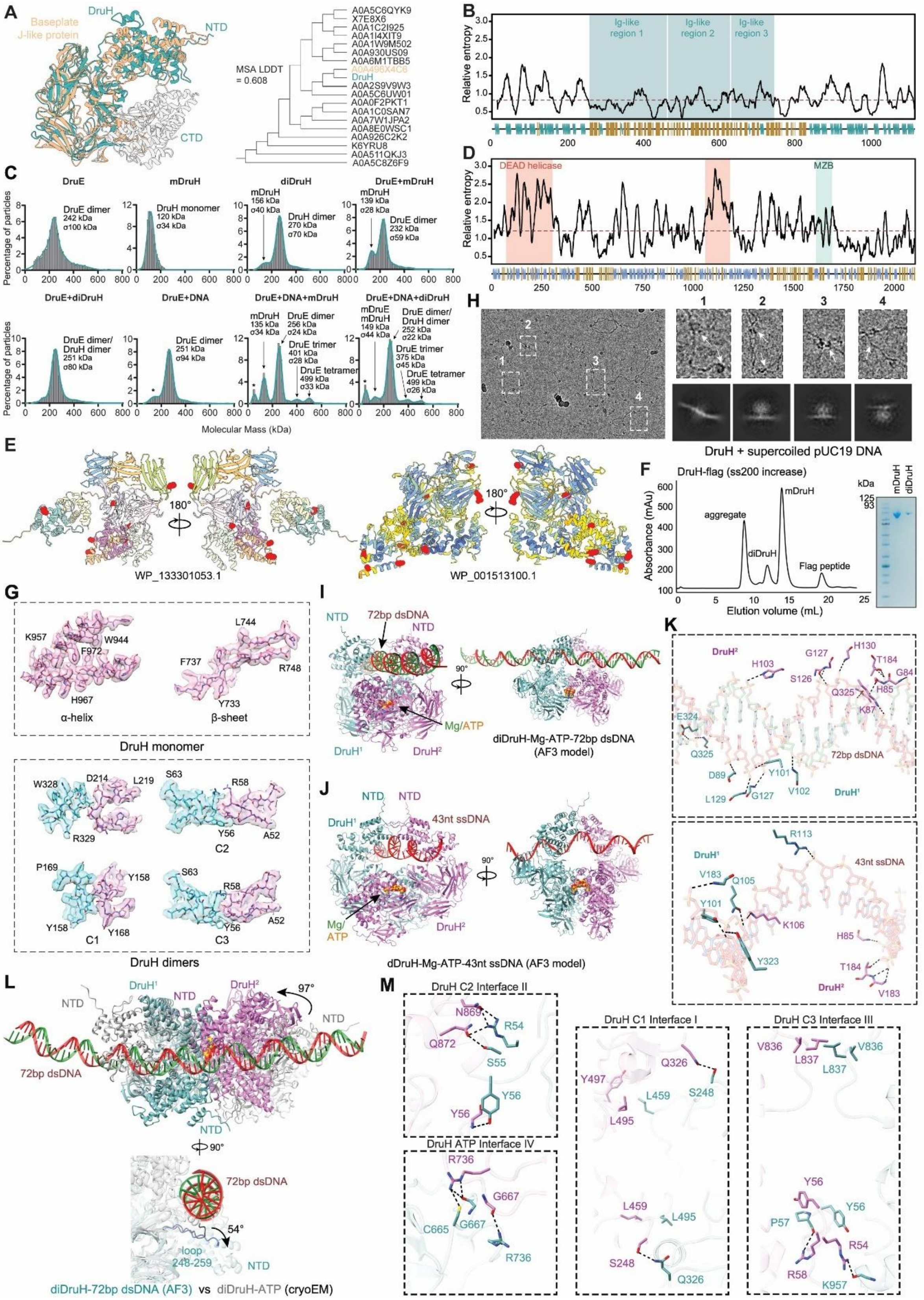


**Figure S6.** Additional evolutionary, biophysical, and structural analysis of DruH. **(A)** FoldMason structural alignment of DruH with representative baseplate J-like proteins recovered by Foldseek. Left, structural superposition of DruH and a representative J-like hit (A0A496X4C6). Right, guide tree of the aligned proteins. The overall MSA LDDT score is indicated. **(B)** Relative entropy across a multiple sequence alignment of 577 representative DruH-family proteins. Ig-like regions, predicted based on structure comparison, are indicated. Red line indicates median relative entropy. **(C)** Representative mass photometry distributions for DruE, monomeric DruH (mDruH), dimeric DruH (diDruH), and the indicated protein mixtures in the absence or presence of DNA. Major fitted species are indicated above each distribution. **(D)** Relative entropy across a multiple sequence alignment of 555 representative DruE-family proteins. DEAD helicase and MrfA Zn-binding (DUF1998) domains are indicated. Red line indicates median relative entropy. **(E)** Residues under positive selection in DruH proteins corresponding to DruE from cluster 11 (WP_020219138.1, left) and cluster 2 (WP_000213430.1, right), mapped in red onto AlphaFold 3 (AF3) models of DruH WP_133301053.1 (left) or WP_001513100.1 (right). Left views are coloured according to TED domain classification, with light pink, orange, blue, green, and violet indicating Ig-like folds. Right views are coloured according to pLDDT confidence scores. **(F)** Size-exclusion chromatography profile of DruH-Flag on a Superdex 200 Increase column, showing aggregate, dimeric DruH (diDruH), monomeric DruH (mDruH), and free Flag peptide. The corresponding Coomassie-stained SDS-PAGE gel of the monomeric and dimeric peak fractions is shown on the right. **(G)** Close-up views of representative local structural features of DruH monomer and dimer conformations, including α-helical and β-sheet elements in the monomer and three representative regions in the dimer conformations (C1-C3). **(H)** Cryo-EM micrograph of DruH incubated with supercoiled pUC19 DNA, with representative particles highlighted. Representative 2D class averages are shown below. **(I)** AF3-predicted model of the DruH dimer bound to 72-bp dsDNA in the presence of $Mg^{2+}$ and ATP, shown in two views. **(J)** AF3-predicted model of the DruH dimer bound to 43-nt ssDNA in the presence of $Mg^{2+}$ and ATP, shown in two views. **(K)** Close-up views of residues predicted to contact the DNA substrates in the AF3 models shown in (I) and (J). **(L)** Superposition of the AF3-predicted DruH dimer bound to 72-bp dsDNA with the cryo-EM structure of the ATP-bound DruH dimer, showing repositioning of the N-terminal domain (NTD). The right panel highlights the rotation angles and the loop spanning residues 248–259. **(M)** Close-up views of residues contributing to the four DruH dimer interfaces identified across the cryo-EM structures: interface I in diDruH-c1, interfaces II and III in diDruH-c2/c3, and interface IV in the ATP-bound dimer.

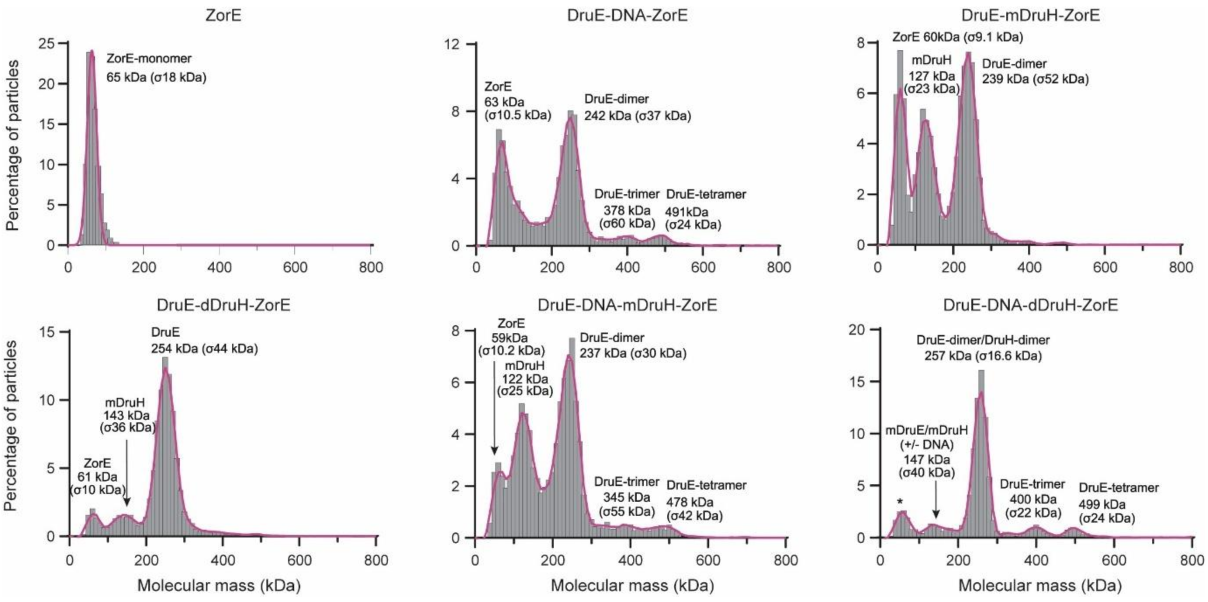


**Figure S7.** Additional analyses of ZorE assemblies. Mass photometry distributions for ZorE alone and for the indicated mixtures containing DruE, DruH, ZorE, and DNA. Major fitted species are indicated above each distribution. Asterisk indicates peaks present in the buffer.
